# Cytoplasmic contractile injection systems mediate cell death in *Streptomyces*

**DOI:** 10.1101/2022.08.09.503279

**Authors:** Bastien Casu, Joseph W. Sallmen, Susan Schlimpert, Martin Pilhofer

**Affiliations:** Department of Biology, Institute of Molecular Biology & Biophysics, Eidgenössische Technische Hochschule Zürich, Otto-Stern-Weg 5, 8093 Zürich, Switzerland; John Innes Center, Department of Molecular Microbiology, Norwich Research Park, Norwich, NR4 7UH, United Kingdom

## Abstract

Contractile injection systems (CISs) are bacteriophage tail-like structures that mediate bacterial cell-cell interactions. While CISs are highly abundant across diverse bacterial phyla, representative gene clusters in Gram-positive organisms remain poorly studied.

Here we characterize a CIS in the Gram-positive multicellular model organism *Streptomyces coelicolor* and show, that in contrast to most other CISs, *S. coelicolor* CIS (CIS^Sc^) mediate cell death in response to stress and impact cellular development. CIS^Sc^ are expressed in the cytoplasm of vegetative hyphae and not released into the medium. Our cryo-electron microscopy structure enabled the engineering of non-contractile and fluorescently tagged CIS^Sc^ assemblies. Cryo-electron tomography showed that CIS^Sc^ contraction is linked to reduced cellular integrity. Fluorescence light microscopy furthermore revealed that CIS^Sc^ contraction mediates cell death upon encountering different types of stress. Finally, the absence of functional CIS^Sc^ had an impact on hyphal differentiation and secondary metabolite production.

Our results provide new functional insights into CISs in Gram-positive organisms and a framework for studying novel intracellular roles, including regulated cell death and life cycle progression in multicellular bacteria.

## Introduction

Bacteria exist in highly competitive environments that require them to interact with a range of organisms. To respond to potential stressors, bacteria have evolved complex strategies to mediate potential antagonistic interactions^1^. One such response is the deployment of cell-puncturing nanodevices called bacterial contractile injection systems (CIS), which are large macromolecular protein machines that can translocate cytotoxic effectors into the extracellular space or directly into target cells^2–5^. In general, CIS are composed of a contractile sheath that encloses an inner tube loaded with effectors, which is fitted with a baseplate complex that facilitates attachment to the membrane. A conformational change in the baseplate complex triggers the contraction of the outer sheath, which leads to the propulsion of the inner tube into the target.

Phylogenetic analyses have indicated that these CISs are conserved across diverse microbial phyla including Gram-negative and Gram-positive bacteria, as well as archaea^6, 7^. CIS are commonly classified as Type VI secretion systems (T6SS) or extracellular CIS (eCIS), based on their modes of action. Anchored at the host’s cytoplasmic membrane, T6SSs function by a cell-cell contact-dependent mechanism, wherein the T6SS injects effectors directly into a neighboring cell^8–12^. By contrast, eCIS are assembled in the bacterial cytoplasm of the donor cell and are subsequently released into the extracellular space, where they can bind to the surface of a target cell, contract and puncture the cell envelope^13^. Recently, a third mode of action was described in multicellular *Cyanobacteria*^14^. This system is also assembled in the bacterial cytoplasm and it then attaches to the thylakoid membrane where it potentially induces lysis of the cell upon stress, resulting in the formation of “ghost cells” which may in turn proceed to interact with other organisms^14^.

Of the hundreds of putative CIS gene clusters detected in sequenced bacteria, all well characterized examples have come from two closely related clades and have been exclusively examined in Gram-negative bacteria. Characterized CIS representatives include “metamorphosis-associated contractile structures” (MACs) from *Pseudoalteromonas luteoviolacea*^15^, the “T6SS subtype *iv*” (T6SS*^iv^*) in *Candidatus* Amoebophilus asiaticus^16^, “antifeeding prophages” (AFPs) from *Serratia*^17^, “Photorhabdus Virulence Cassettes” (PVCs) from *P. asymbiotica*^18^, and two newly characterized CIS from the marine bacteria *Algoriphagus machipongonensis*^19^ and *Cyanobacteria*^14^.

Strikingly, 94 of 116 sequenced Gram-positive actinomycetes of the genus *Streptomyces* were shown to encode a potential CIS gene cluster^6, 7^. A previous report suggested that CIS from *S. lividans* were involved in microbial competition, however, the mechanism remains unknown^20^. *Streptomyces species* are abundant soil bacteria and renowned for their complex developmental life cycle and their ability to produce an array of clinically relevant secondary metabolites^21^. The *Streptomyces* life cycle begins with the germination of a spore and the generation of germ tubes which grow by apical tip extension and hyphal branching to form a dense vegetative mycelium. Upon nutrient depletion, non-branching aerial hyphae are erected, which eventually synchronously divide into chains of uni-nucleoid spores^22^. Notably, the production of these important molecules is tightly coordinated with the developmental life cycle^21^.

Here, we provide evidence that CISs from the model organism *Streptomyces coelicolor* (CIS^Sc^) function intracellularly and belong to a new class of contractile injection systems that exist as free-floating, fully assembled particles in the cytoplasm and mediate cell death in response to stress conditions. Additionally, we found that the absence of CISs affects the coordinated cellular development and secondary metabolite production of *S. coelicolor*, indicating a wider role of CIS from *Streptomyces* in the multicellular biology of these important bacteria.

## Results and Discussion

### *Streptomyces* express cytoplasmic CIS during vegetative growth

Previous bioinformatic studies revealed that the majority of sequenced *Streptomyces* genomes harbor a highly conserved cluster of eCIS genes related to the poorly studied CIS IId subtype^6, 7^. This was further confirmed by our phylogenetic analyses using sheath protein sequences from known producers of CIS and from two representative *Streptomyces* species, namely *S. coelicolor* and *S. venezuelae* (Fig. 1a).

**Figure 1:**
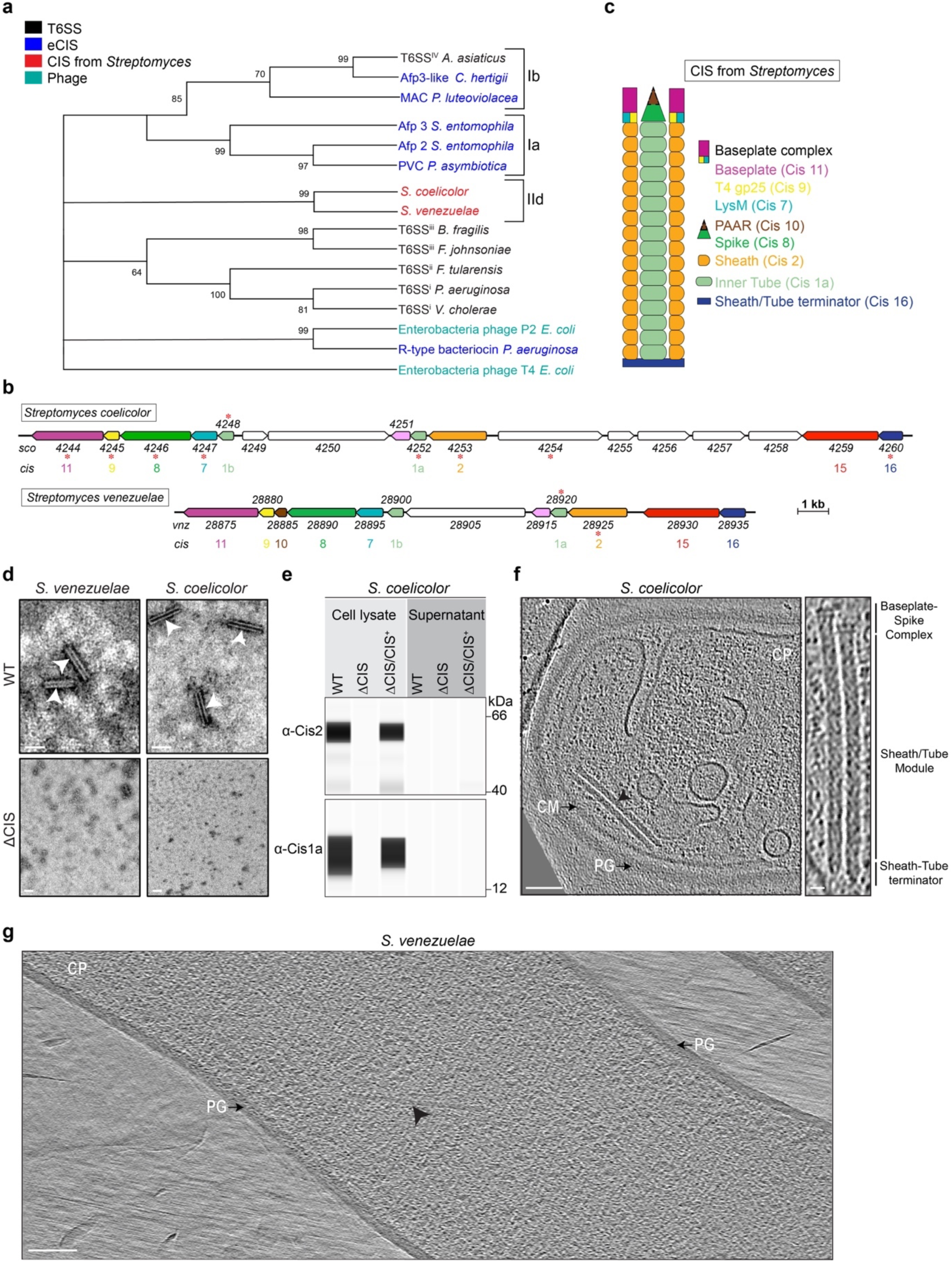
Different *Streptomyces* species express cytoplasmic CIS assemblies. **a.** Phylogenetic analysis of representative sheath protein sequences shows that homologs from *Streptomyces* form a monophyletic clade. Numbers indicate bootstrap values, color code denotes different modes of action. Subclades Ia, Ib and IId are based on the dbeCIS database^6^. **b.** Representative gene clusters from *Streptomyces* encode conserved CIS components. The schematic shows the gene arrangement of the CIS gene clusters from *S. coelicolor A3(2)* (CIS^Sc^) and *S. venezuelae NRRL B-65442* (CIS^Sv^) with gene locus tags. Color code indicates conserved gene products. CIS components were numbered based on similarities to previously studied CIS (AFP)^14, 63^. Asterisks indicate gene products that were detected by mass spectrometry after CISs purification (Supplementary Table 1). **c.** The schematic illustrates a putative CIS assembly from *Streptomyces*. Color-code is based on the predicted gene function shown in (b). **d.** The gene *cis2* is required for CIS assembly. Shown are negative-stain EM images of crude sheath preparations from WT and 11CIS mutant strains of *S. coelicolor* and *S. venezuelae*. White arrowheads indicate contracted sheath-like structures. Shown are representative micrographs of three independent experiments. Bars, 80 nm. **e.** CIS^Sc^ proteins are detected in the cell lysate but not secreted into the supernatant. Shown is the automated Western blot analysis of cultures of *S. coelicolor* WT, 11CIS mutant, and a complementation (11CIS/CIS^+^). The presence of the sheath protein (Cis2) and the inner tube protein (Cis1a) in whole cell lysates and concentrated culture supernatants was probed using polyclonal antibodies against Cis1a/2. Experiments were performed in biological replicates. For the control SDS-PAGE gel see Extended Data Fig. 1. **f.** Shown is a cryo-electron tomogram of a WT *S. coelicolor* hypha, revealing two cytoplasmic extended CIS^Sc^ assemblies (arrowhead). PG, peptidoglycan; CM, cytoplasmic membrane; CP, cytoplasm. Putative structural components are indicated on the right. Bars, 75 nm and 12.5 nm (magnified inset). **g.** Shown is a cryo-electron tomogram of a cryoFIB milled WT *S. venezuelae* hypha, revealing one cytoplasmic extended CIS^Sv^ assembly (arrowhead). PG, peptidoglycan; CP, cytoplasm. Bar, 140 nm.

Closer inspection of the *Streptomyces* CIS gene clusters from *S. coelicolor* (*sco4244-Sco4260*) and *S. venezuelae* (*vnz_28875-vnz_28935*) suggested that both species encode 10 and 11 core structural components of the phage-tail-like systems, respectively^6, 7^ (Fig. 1b/c). Based on this sequence similarity, we renamed the genes from *Streptomyces* to *cis1-16*. Both CIS gene clusters encode two inner tube homologs (*cis1a* and *cis1b*) as well as additional proteins of unknown function. Cis10, a PAAR-repeat containing protein, is only present in *S. venezuelae*. Genes encoding a tail fiber protein (Afp13), which mediates eCIS binding to target cells, and a tail measure protein (Afp14), involved in regulating the length of eCIS particles^17^, are absent in both CIS gene clusters.

To test whether *S. coelicolor* and *S. venezuelae* produced CIS, we purified sheath particles from crude cell lysates, followed by negative stain electron microscopy (EM) imaging. We observed typical contracted sheath-like particles in crude extracts from wild-type (WT) *S. coelicolor* and *S. venezuelae*, while no such assemblies were seen in strains carrying a deletion in *cis2* (ΔCIS, Fig. 1d). Subsequent mass spectrometry analysis of the purified particles detected peptides from Cis1a (inner tube) and Cis2 (sheath) (Extended Data Table 1), confirming that the CIS gene clusters from *Streptomyces* encode CIS-like complexes. We noticed that *S. coelicolor* produced approximately 50 times more sheath particles compared to *S. venezuelae* (Extended Data Fig. 1a/b). Therefore, we focused on the characterization of CIS from *S. coelicolor* (CIS^Sc^) in subsequent experiments.

To test if CIS^Sc^ displayed a mode of action similar to canonical eCIS, we investigated whether CIS^Sc^ were released from cells into the extracellular space. Using automated Western blotting, we analyzed the culture supernatant and whole cell extracts from WT and ΔCIS *S. coelicolor* cells that were grown for 48 h in liquid medium. Interestingly, we detected the two key CIS^Sc^ components Cis1a (inner tube) and Cis2 (sheath) only in whole cell lysates but not in the supernatant of cultures of the WT or the complemented ι1CIS mutant (Fig. 1e). SDS-PAGE analysis of concentrated culture supernatants further confirmed that all tested samples contained protein (Extended Data Fig. 1c). These findings suggest that the entire CIS^Sc^ assembly is retained in the cytoplasm, unlike typical T6SS (inner tube protein translocated into the medium) and unlike eCIS (full assemblies released into the medium)^8, 19^. Next, to visualize the localization of CIS^Sc^ *in situ*, we imaged hyphae of *S. coelicolor* and *S. venezuelae* by cryo-electron tomography (cryoET). While intact *S. coelicolor* hyphae could be imaged directly, *S. venezuelae* was too thick and had to be thinned by cryo-focused ion beam (FIB) milling prior to imaging. We predominantly found extended CIS that appeared to be free-floating in the cytoplasm, a behavior that is inconsistent with a T6SS mode of action (Fig. 1f/g). Taken together, these results indicate that CIS from *Streptomyces* may play a role in intracellular processes, which would be distinct from the previously described functions for T6SS and eCIS.

### Structure, engineering and subcellular localization of CIS^Sc^

To obtain insights into the structural details of the CIS^Sc^ contractile sheath-tube module, we performed single particle cryoEM (helical reconstruction) of purified sheath particles from WT *S. coelicolor*, which had a homogeneous length of ∼140 nm (Fig. 2a-c). The resulting map of the contracted sheath reached a resolution of 3.6 Å (Extended Data Fig. 2a/b). Contracted sheath proteins adopt a right-handed helical array with an inner diameter of 115 Å and an outer diameter of 233 Å (Fig. 2b). Similar to the recently described sheath structures observed in AlgoCIS^19^ and tCIS^14^, the CIS^Sc^ sheath is comprised of only one protein (Cis2). Cis2 monomers consist of three domains and are well conserved in *S. coelicolor* and *S. venezuelae*, sharing ∼65% sequence identity (Extended Data Fig. 2c). From the resulting map, it was possible to build *de novo* domains 1 and 2, which contribute to the sheath wall (Fig. 2c). The additional domain 3, which is located on the sheath surface, seems to be highly flexible. The overall contracted structure of Cis2 is similar to sheaths of previously characterized systems^18, 23, 24^.

**Figure 2:**
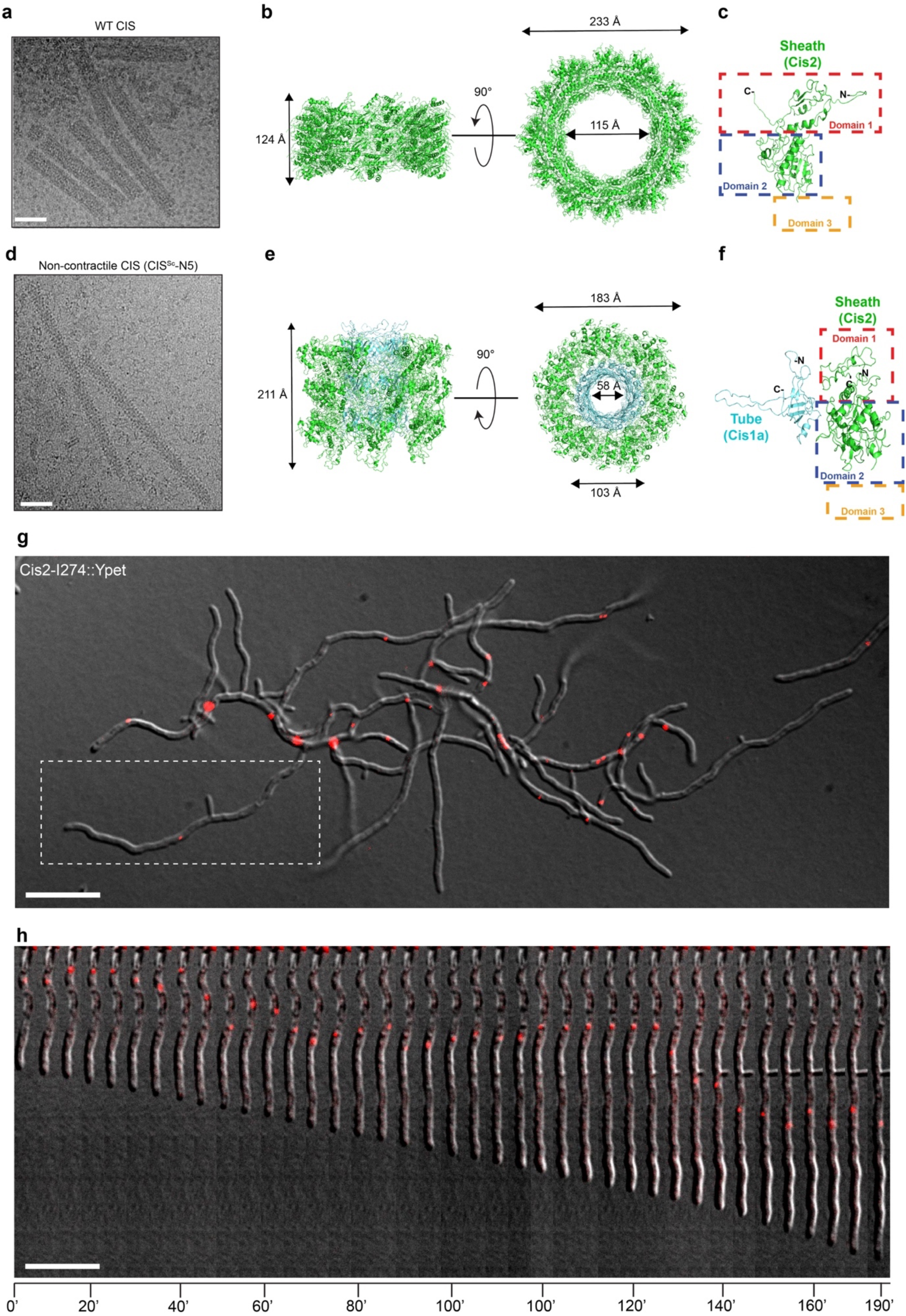
Structure and subcellular localization of CIS^Sc^. **a.** Shown is a representative cryo-electron micrograph of a sheath preparation from WT *S. coelicolor* that was recorded for structure determination. All sheath structures were seen in the contracted state. Bar, 40 nm. **b.** Shown is a section of the CIS^Sc^ sheath cryoEM structure in the contracted conformation. **c.** Shown is a ribbon representation of the Cis2 monomer in its contracted state. Dashed rectangles highlight the positions of domains 1 (red), 2 (blue) and 3 (orange, not resolved because of high flexibility). **d.** Shown is a representative cryo-electron micrograph of a sheath preparation from *S. coelicolor* expressing a non-contractile mutant of Cis2 (CIS-N5). More than 95% of all structures were seen in the extended state. Bar, 40 nm. **e.** Shown is ribbon representation of a section of the *S. coelicolor* Cis2 (sheath)-Cis1a (inner tube) cryoEM structure in the extended conformation that was solved using the non-contractile mutant. **f.** Shown is a ribbon representation of the Cis2 monomer (non-contractile mutant) in its extended state. Dashed rectangles highlight the positions of domains 1 (red), 2 (blue) and 3 (orange, not resolved because of high flexibility). **g.** Insights from the cryoEM structures enabled us to tag Cis2 with a fluorescent tag (YPet) for subsequent time-lapse imaging to determine the localization of assembled CIS^Sc^. Shown is a still image from Supplementary Movie 1, showing scattered fluorescent foci inside vegetative hyphae. Cells were first grown in TSB-YEME for 40 h and then spotted onto an agarose pad prepared from culture medium and subsequently imaged by time-lapse fLM. White rectangle highlights hypha shown in (h). Bar, 10 µm. **h.** Fluorescently tagged CIS^Sc^ remained largely static or showed short-range movements over time. Shown is an image montage of a representative growing *S. coelicolor* hypha from Supplementary Movie 1. Images were acquired every 5 min. Bar, 10 µm.

In order to be able to purify the extended form of the CIS^Sc^ sheath tube module from *S. coelicolor* cell lysates, we set out to engineer non-contractile CIS^Sc^ based on the information from the contracted Cis2 structure and based on similar previous approaches in *V. cholerae*^25^ and enteroaggregative *E. coli*^26^ (Extended Data Fig. 3a). Different sets of two (IE), three (IEG) and five (IEGVG) amino acid residues were inserted into the N-terminal linker of Cis2 after position G25, resulting in the mutants CIS-N2, CIS-N3, and CIS-N5, respectively. For the CIS-N2 and CIS-N3 mutants, less than 30% and 50% were found in extended form, respectively (Extended Data Fig. 3b/c). For the CIS-N5 non-contractile mutant, more than 95% of the complexes were seen in the extended conformation (Extended Data Fig. 3d). *In vitro*, the length of the CIS-N5 non-contractile mutant was homogeneous at ∼230 nm (Fig. 2d). Moreover, mass spectrometry analyses confirmed the presence of most CIS^Sc^ components, indicating the stability of the complex (Fig. 1b, Extended Data Table 1).

Next, we optimized the purification of CIS-N5 and performed cryoEM. Helical reconstruction was used to generate an EM map, which we then used to build *de novo* the sheath-tube (Cis2-Cis1a) module in extended conformation at 3.9 Å resolution (Extended Data Fig. 3e/f). Domain 3 of the extended sheath (Cis2) was again too flexible to be resolved. The tube (Cis1a) structure and fold are highly similar to the tube structures already described for other CISs (Fig. 2f, Extended Data Fig. 3g). The comparison of the sheath (domains 1/2) in the extended vs. the contracted states revealed an increase in diameter and shortening of the length upon contraction, similar to other CISs (Fig. 2b/e)^14, 18, 19, 24, 27^.

Guided by the high-resolution structure of the sheath module (Fig. 2a-f), we engineered a fluorescently tagged CIS^Sc^ by inserting YPet at position I274 in the Cis2 monomer. Subsequently, we used this Cis2-YPet sandwich fusion to complement the *S. coelicolor* 11*cis2* mutant *in trans* (Extended Data Fig. 4a). Using negative stain EM and cryoET, we confirmed that YPet-tagged CIS^Sc^ were able to assemble into extended particles and to contract, suggesting that these fluorescently labeled CIS^Sc^ particles were functional (Extended Data Fig. 4b/c). This enabled us to visualize the subcellular localization of CIS^Sc^ in vegetatively growing hyphae using time-lapse fluorescence light microscopy (fLM). Multiple CIS^Sc^-YPet foci were found inside the hyphae but not in extracellular space. The foci were largely static or displayed short-range movements within the hyphae (Fig. 2g/h and Extended Data Movie 1). CIS^Sc^-YPet foci were stable over the course of the experiment and did not reveal significant changes in the shape or intensity of the fluorescence. While this indicates the absence of firing events during the experiment, the resolution in fLM and the relatively short length of the CIS^Sc^ may hamper the detection of firing events (in contrast to the much longer T6SSs^8, 28^).

Taken together, our structural data allowed us to engineer non-contractile and fluorescently tagged CIS^Sc^, which revealed the presence of scattered CIS^Sc^ in *S. coelicolor* hyphae.

### CIS^Sc^ contraction state correlates with the integrity of the cell membrane

Our initial cryoET data of *S. coelicolor* cells indicated that contracted CIS^Sc^ were frequently found in hyphae that displayed a damaged cell membrane. To explore this correlation further, we first acquired low magnification two-dimensional (2D) cryoEM images. Based on the contrast of individual hyphae in these 2D images (Fig. 3a), we classified the hyphae into three distinct groups: (1) ‘intact hyphae’ (dark appearance in 2D) with mostly intact cytoplasmic membrane and occasional vesicular membranous assemblies that are reminiscent of “cross-membranes”^29^ (Fig. 3b); (2) ‘partially lysed hyphae’ with a mostly disrupted/vesiculated cytoplasmic membrane (reduced contrast in 2D), indicative of cytoplasmic leakage (Fig. 3c); and (3) membrane-less ‘ghost cells’ (lysed hyphae; hardly visible in 2D) that only consisted of the peptidoglycan cell wall (Fig. 3e). Representative hyphae of each group (n=90) were imaged by cryoET (270 tomograms in total, n=3 experiments) and the conformational state and *in situ* localization of the CIS^Sc^ was determined (Fig 3b-g). In addition, we performed 3D volume segmentation of selected full tomograms.

**Figure 3:**
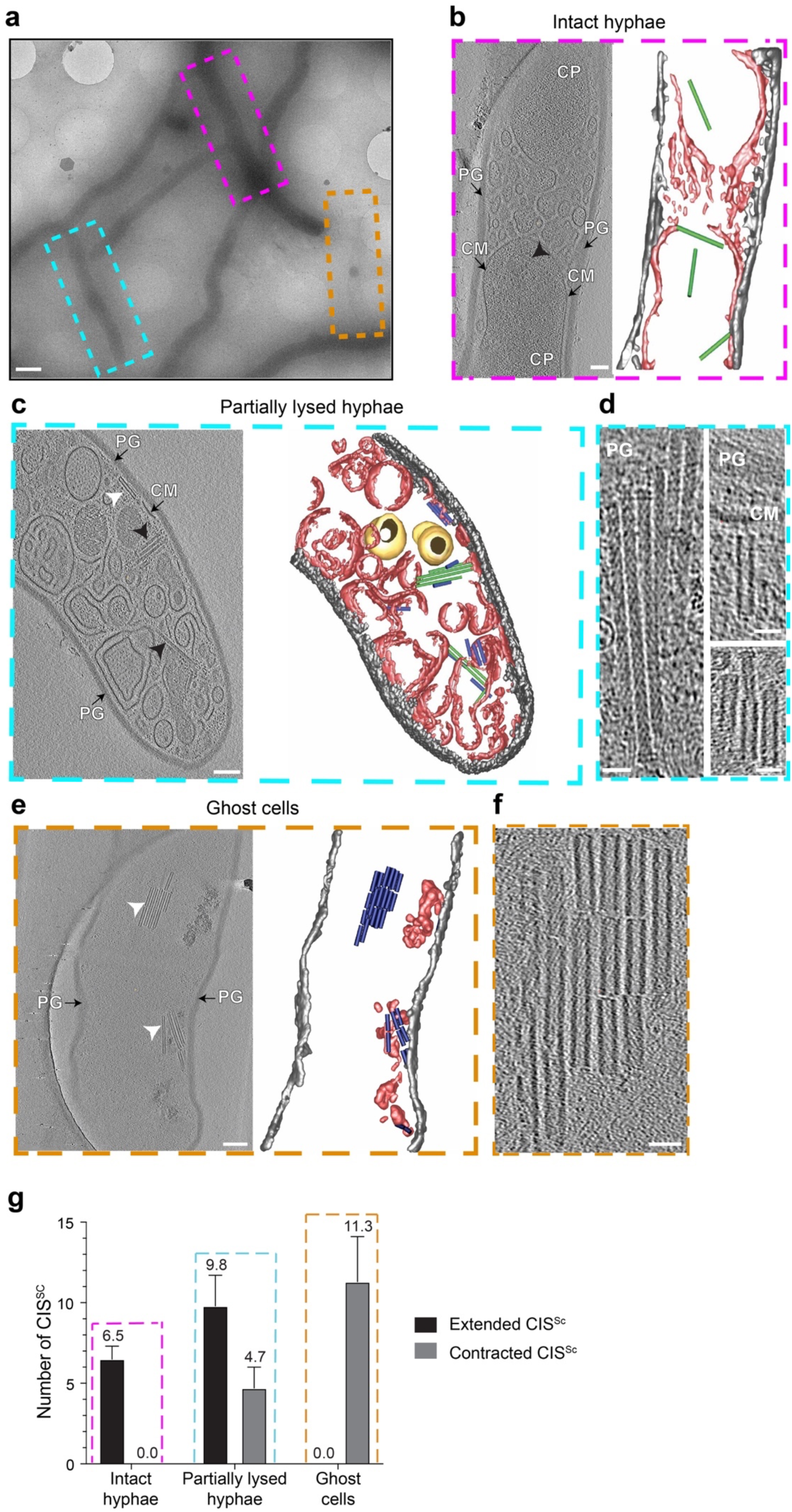
Sheath contraction is linked to reduced cellular integrity. **a.** Shown is a representative low magnification 2D cryoEM image of WT *S. coelicolor* hyphae during vegetative growth. Hyphae were divided into three classes based on their density in such images and based on their structure in cryo-tomograms: (1) ‘intact hyphae’ (purple box), (2) ‘partially lysed hyphae’ (cyan box), and (3) ‘ghost cells’ (orange box). Bar, 1 μm. **b-f.** Shown are representative cryo-tomographic slices and 3D renderings of hyphae of the three classes (corresponding to the regions boxed in a). ‘Intact hyphae’ (b) had a mostly intact cytoplasmic membranes and occasional vesicular membranous assemblies that are reminiscent of “cross-membranes”^29^. ‘Partially lysed hyphae’ (c) showed a mostly disrupted/vesiculated cytoplasmic membrane. ‘Ghost cells’ (e) contained only remnants of membranes and a mostly intact peptidoglycan cell wall. Note the frequent occurrence of CIS^Sc^ assemblies in extended (black arrowheads/green) and contracted (white arrowheads/blue) conformations. Magnified views of clusters of CIS^Sc^ seen in cryo-tomograms are shown in d/f. PG/grey, peptidoglycan; CM/red, cytoplasmic membrane/membranes; CP, cytoplasm; yellow, storage granules. Bars, 75 nm in b/c/e and 25 nm in d/f. **g.** Sheath contraction correlates with cellular integrity, showing the presence of only extended CIS^Sc^ in the class ‘intact hyphae’, and the presence of only contracted CIS^Sc^ in ‘ghost cells’. Shown is a quantification of extended and contracted CIS^Sc^ per tomogram of WT *S. coelicolor* hyphae. Results are based on three independent experiments, with n=30 tomograms for each class of cells.

As observed before for intact hyphae (Fig. 1f), individual CIS^Sc^ particles were always found in the extended conformation and localized in the cytoplasm (Fig. 3b). By contrast, in partially lysed hyphae (Fig. 3c), the ratio of extended to contracted CIS^Sc^ was 2:1. CIS^Sc^ particles often appeared to cluster in the vicinity of membranous structures (Fig. 3d). Notably, we found that in some cases, the extended CIS^Sc^ aligned perpendicular to membrane patches or vesicles with the baseplate complex facing the membrane, indicating that CIS^Sc^ may interact with the cytoplasmic membrane (Fig. 3c/d). In contrast, ghost cells only displayed CIS^Sc^ particles in the contracted state and which were often clustered (Fig. 3f).

Collectively, these results indicated that the conformational state of CIS^Sc^ correlates with the integrity of the cell and that CIS^Sc^ may play an intracellular role as a consequence of cellular stress and either directly or indirectly lead to cell death. Hence, we hypothesized that such stress conditions could result in the recruitment of CIS^Sc^ to the membrane and trigger firing.

### CIS^Sc^ c**o**ntraction mediates cell death under stress conditions

To test this hypothesis, we explored whether upon encountering stress, the presence of CIS^Sc^ and their contraction could mediate cell death. To generate a marker for cell viability, we inserted *sfgfp* under the control of a constitutive promoter *in trans* in *S. coelicolor* WT, in the 11CIS null mutant, and in the non-contractile mutant (CIS-N5). In order to label intact and partially lysed hyphae, cells were incubated with the fluorescent membrane dye FM5-95. We first used correlated cryo-light and electron microscopy (CLEM) to confirm that the detected cytoplasmic and membrane fluorescence correlated with the physiological state of the hyphae (Extended Data Fig. 5). To assess the level of cell death in the imaged strains, we used fLM and quantified the ratio of sfGFP signal (indicator of viable hyphae) to FM5-95 signal (indicator of intact and lysed hyphae) in the different strains. Cells were grown for 48 h in liquid, a time-point at which CIS^Sc^ can be detected in hyphae (Fig. 1d/e).

During non-stress conditions, the WT, the ΔCIS and the CIS-N5 mutant strains displayed a similar sfGFP/FM5-95 ratio, indicating that none of the strains showed a significant difference in viability (Fig. 4a/c). In parallel, we challenged the same *S. coelicolor* strains with a sub-lethal concentration of the bacteriocin nisin (1 µg/ml) for 90 min, which causes the formation of membrane pores and eventually will lead to the disruption of cell envelope integrity^30^. In the WT, we found that ∼50% of the analyzed hyphae displayed signs of cell death (Fig. 4b/d). Strikingly, in the CIS-deficient strain and the non-contractile CIS^Sc^ mutant there was no dramatic induction of cell death upon nisin treatment (Fig. 4b/d).

**Figure 4:**
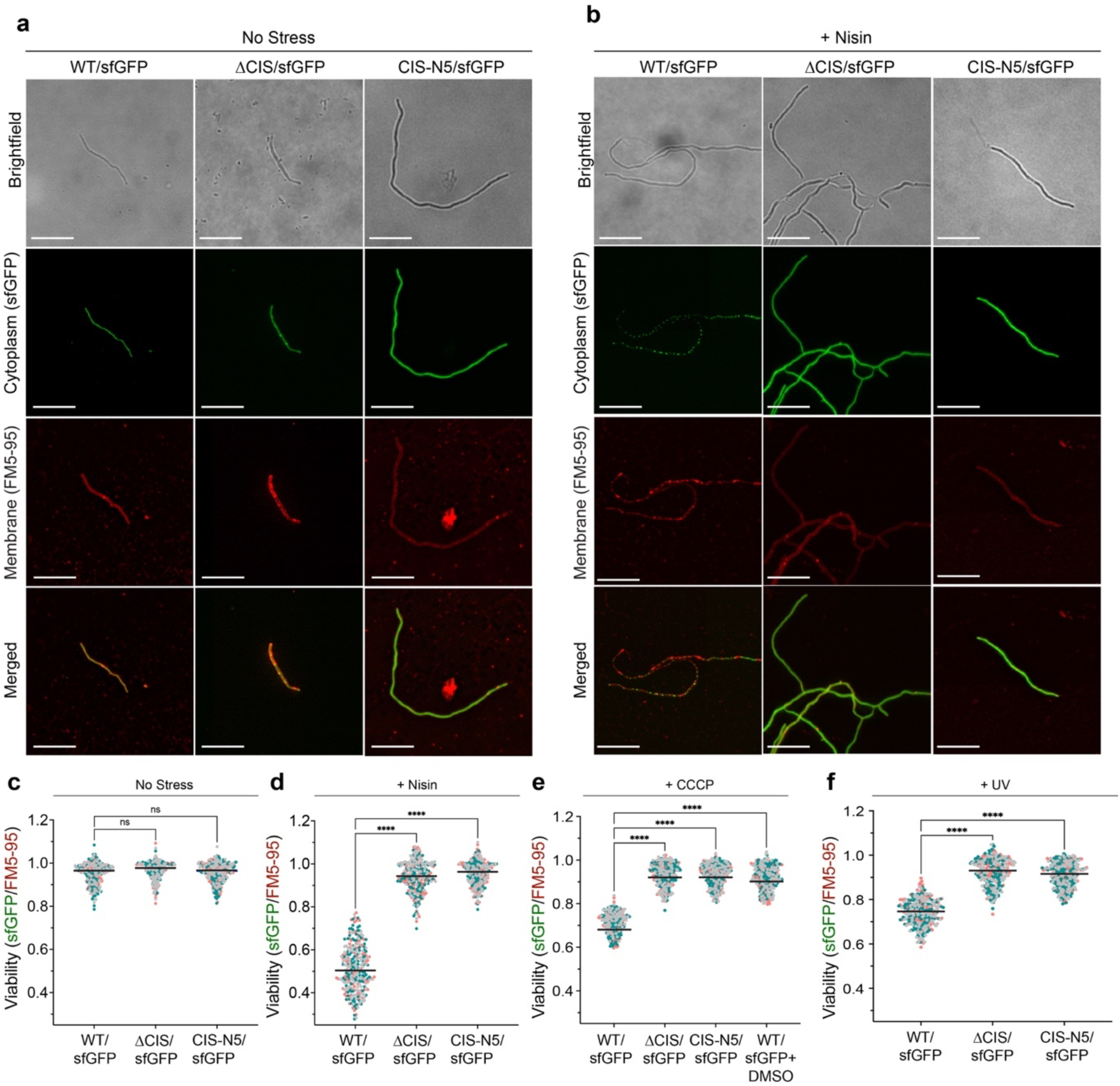
*S. coelicolor* with functional CIS^Sc^ show increased cell death upon stress. **a/b.** fLM (shown are representative images) was used to determine the ratio between live cells (cytoplasmic sfGFP) and total cells (membrane dye FM5-95) after growth in the absence of stress (a) or in the presence of nisin stress (b). *S. coelicolor* WT/sfGFP, ΔCIS/sfGFP and CIS-N5/sfGFP were grown in TSB for 48 h and were then treated with 1 µg/ml nisin for 90 min. Bars, 10 µm. **c/d.** The quantification of the experiments in a/b showed no significant differences between the WT strain and both CIS^Sc^ mutants under conditions without stress. In contrast, nisin-stressed WT cells showed a significantly higher rate of cell death compared to both nisin-stressed mutants. Superplots show the area ratio of live to total hyphae. Black line indicates the mean ratio derived from biological triplicate experiments (n=100 images for each experiment). Ns (not significant) and **** (p < 0.0001) were determined using a one-way ANOVA and Tukey’s post-test. **e/f.** To test the induction of cell death under other stress conditions, the same strains were treated with the protonophore CCCP (10 µM, or 0.002 % DMSO as mock control) (e) or UV light (f) for 10 min. Similar to nisin stress, we detected a significant difference in cell death induction between WT and both CIS^Sc^ mutants. See Extended Data Fig. 6a/b for representative fLM images.

To investigate whether other stress factors could induce cell death, we repeated the experiments and challenged *S. coelicolor* with the membrane depolarising agent CCCP (carbonyl cyanide 3-chlorophenylhydrazone) and with UV stress, to induce DNA damage (Extended Data Fig. 6a/b). In line with our previous results, the treatment of vegetative hyphae with CCCP or with UV radiation both led to an increase in cell death by 25% in the WT but not in hyphae of the ΔCIS or the CIS-N5 mutant strain (Fig. 4e/f). In parallel, we also purified CIS^Sc^ from crude cell extracts obtained from non-stressed and stressed samples that were used for fLM imaging. By negative-stain EM imaging, we confirmed the presence of CIS^Sc^ particles in hyphae of the WT and in the CIS-N5 mutant strain, and the absence of sheath particles in the ΔCIS mutant (Extended Data Fig. 7). The abundance of CIS^Sc^ in non-stressed and stressed samples was comparable, which was also confirmed by the detection of Cis1a/2 proteins by Western blotting analysis of non-stressed vs. nisin-treated hyphae (Extended Data Fig. 8a/b).

### CIS^Sc^ contribute to the multicellular development of *Streptomyces*

Earlier studies indicated that the expression of the *S. coelicolor* CIS gene cluster is coordinated with the *Streptomyces* life cycle^31^. To follow the expression of CIS^Sc^ during the developmental life cycle, we constructed a fluorescent reporter strain in which expression of *ypet* was driven by the *cis2* promoter (*P_cis2_-ypet*). Since *S. coelicolor* only completes its spore-to-spore life cycle when grown on solid medium, glass coverslips were inserted at a 45-degree angle into agar plates inoculated with spores. Coverslips with attached *S. coelicolor* hyphae were removed and imaged every 24 h for four days by fLM. Fluorescent signal indicated that the *cis2* promoter was primarily active in vegetative hyphae at the 48-h time point (Fig. 5a). In parallel, we determined CIS^Sc^ protein levels in surface-grown WT *S. coelicolor* over the life cycle. Consistent with our fluorescence reporter experiment, Cis1a/2 levels were highest in vegetative mycelium that was harvested after 30 h and 48 h of incubation (Extended Data Fig. 9a). These results are also in agreement with published transcriptomics data from *S. venezuelae*, showing the specific induction of the CIS^Sv^ gene cluster during vegetative growth (Extended Data Fig. S9b)^32^.

**Figure 5:**
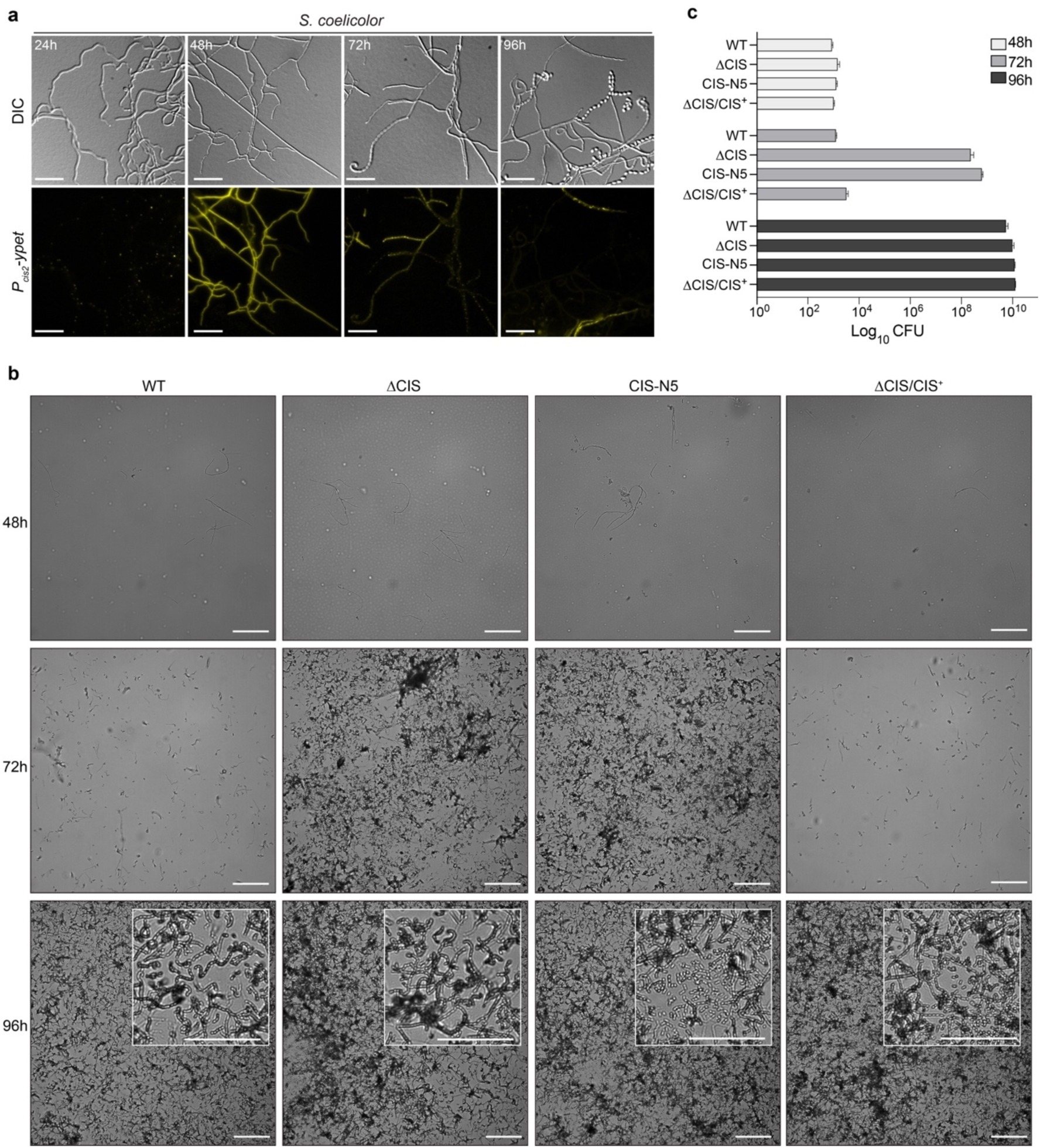
Functional CIS^Sc^ are involved in *Streptomyces* multicellular development. **a.** Microscopic analysis of *S. coelicolor* WT cells expressing a fluorescent promoter fusion to the sheath promoter *p_cis2_*-*ypet in trans*, showing that the sheath operon of the CIS^Sc^ cluster is predominantly expressed during vegetative growth (48 h). Shown are representative micrographs of surface-grown *S. coelicolor* hyphae that attached to a microscopic cover glass inserted into the inoculated agar surface at a 45-degree angle. Plates were incubated over 96 h at 30 °C and imaged at the indicated time-points. Experiments were performed in biological triplicates. Bars, 10 µm. **b.** Representative brightfield images of surface imprints of plate-grown colonies of *S. coelicolor* WT, the CIS^Sc^ mutant strains ΔCIS, CIS-N5, and the complemented mutant ΔCIS/CIS^+^. Images were taken at the indicated timepoints. Only hyphae undergoing sporulation or spores will attach to the hydrophobic cover glass surface. Insets show magnified regions of the colony surface containing spores and spore chains. Note that strains with functional CIS sporulate later. Bars, 50 µm. **c.** Shown is a quantification of spore production (colony forming unit, CFU) in the same strains as above, revealing at 72 h much higher CFUs (spores) in both CIS mutants. Strains were grown on R2YE agar and spores were harvested after 48 h, 72 h and 96 h of incubation. Data shows mean values and standard deviation obtained from biological triplicate experiments.

Since a previous study on *S. lividans* reported a putative role of CISs in inter-species interactions^20^, we performed a series of growth competition assays but did not observe any obvious differences in fitness between the WT and CIS^Sc^-mutants (Supplementary Table 2). We therefore then tested whether the expression of functional CIS^Sc^ had an effect on the timely progression of the *S. coelicolor* life cycle, using WT, 11CIS, CIS-N5 and a complemented strain. First, we detected sporulating hyphae and spores by imaging surface imprints of plate grown (R2YE agar) colonies at different time points. All strains consistently completed their life cycle and synthesized spores (Fig. 5b). Importantly, in contrast to the WT and the complemented strain, both ΔCIS and CIS-N5 mutants sporulated markedly earlier (72 h vs. 96 h for the WT and the complemented mutant). These results were further corroborated by quantifying the number of spores produced by the individual strains under the same experimental conditions (Fig. 5c).

In addition to the accelerated cellular development in CIS^Sc^ mutants, we also noticed by the appearance of the cultures from same strains grown in liquid R2YE, that production of the two characteristic pigmented secondary metabolites in *S. coelicolor*, actinorhodin (blue)^33^ and undecylprodigiosin (red)^34^, was significantly reduced, compared to the WT and the complemented 11CIS mutant (Extended Data Fig. 9c). This was further confirmed by a quantification of the total amount of actinorhodin (intracellular and secreted) produced over a period of 72 h. Both ΔCIS and CIS-N5 mutants produced approximately 70 % less actinorhodin compared to the WT and the ΔCIS complementation strain (ΔCIS/CIS^+^) (Extended Data Fig. 9d). Moreover, in contrast to the observed delay in sporulation, the actinorhodin production in the CIS^Sc^ mutants was not just delayed, but it never reached WT levels until the end of the experiment.

Altogether, we showed that deleting or expressing non-functional CIS^Sc^ results in significant changes in the *S. coelicolor* life cycle progression, which also affects secondary metabolite production.

## Conclusions

Here we show that CIS particles from *Streptomyces* are functionally distinct from related eCIS and T6SS. Our data from fLM imaging, cryoET imaging and Western blotting all indicate consistently that CIS^Sc^ were assembled free floating in the cytoplasm, however, under our experimental conditions, they were not found to be released into the medium, nor were they seen attached to the cytoplasmic membrane. This argues against a mode of action as a typical eCIS. In addition the *Streptomyces* CIS gene cluster does not contain a typical tail fiber-like protein for binding of a potential target cell. This also speaks against a typical T6SS mode of action, since it is difficult to imagine how a CIS^Sc^ acting as a T6SS would fire through the thick peptidoglycan cell wall in the Gram-positive host organism. Therefore, our data points to an intracellular function, which is supported by further observations that are discussed below.

CryoET imaging revealed a significant fraction of partially or fully lysed cells in a vegetative culture. Interestingly, the degree of cell lysis strongly correlated with the presence of contracted CIS^Sc^ assemblies. fLM imaging, on the other hand, showed that under different types of stress conditions, cell death was induced in a WT strain but significantly less in mutants that did not express CIS^Sc^ or that expressed non-contractile CIS^Sc^. Importantly, cell viability was not compromised by these stress conditions in CIS-deficient or non-contractile mutant strains. CIS^Sc^ contraction is therefore required for inducing cell death once a culture encounters stress. We speculate that cell lysis could be achieved by membrane- or cell wall-targeting effectors that are loaded into the CIS^Sc^ and released upon contraction. This could either happen upon CIS^Sc^ binding to the cytoplasmic membrane followed by contraction, or by contraction of a free floating CIS^Sc^ releasing effectors into the cytoplasm, which in both examples might trigger cell death. A similar mode of action was recently proposed for thylakoid-anchored CISs in multicellular cyanobacteria^14^.

In addition to mediating death of the host cell in response to stress, we showed that CIS^Sc^ contraction also plays a role in the timely progression of the *Streptomyces* life cycle, evidenced by the earlier onset of sporulation in CIS^Sc^ mutants. Cell death has been proposed as a distinct process in the developmental programme of *Streptomyces*^35^. However, the underlying molecular mechanism has remained unclear. We speculate that contracting CIS^Sc^ could induce hyphal cell death, which impacts the *Streptomyces* multicellular development. Notably, increased cell death has been reported to occur at the center of colonies^36, 37^. These regions are thought to be limited in nutrient and/or oxygen supply, which in turn may be perceived as stress and trigger CIS^Sc^-mediated cell death.

In addition, the morphological differentiation of *Streptomyces* colonies is tightly coordinated with the production of secondary metabolites, which are often secreted into the environment where they can provide a competitive advantage^21^. We showed that CIS^Sc^ mutants were not only significantly affected in the timing of the onset of sporulation, but also in the production of the secondary metabolite actinorhodin. We speculate that the delay of sporulation in the WT (and the complemented strain) may be advantageous to allow the coordinated production and release of key secondary metabolites such as toxins, proteases or signaling molecules. The lack of functional CIS^Sc^ in both mutant strains could lead to improper timing of cell cycle progression, resulting in early sporulation, which may in turn lead to lower amounts of actinorhodin production.

In conclusion, our data provide new functional insights into CISs in a Gram-positive model organism and a framework for studying new intracellular roles of CIS, including regulated cell death and life cycle progression.

## Acknowledgements

We thank ScopeM for instrument access at ETH Zürich. We thank the Functional Genomics Center Zürich for mass spectrometry support. Pilhofer Lab members are acknowledged for discussions. Jingwei Xu and Armin Picenoni are acknowledged for input on SPA cryoEM data collection. We thank Matt Bush for help with the automated Western blot analysis. M.P. was supported by the Swiss National Science Foundation (no. 31003A_179255), the European Research Council (no. 679209), and the NOMIS foundation. Work in the lab of S.S. was supported by a Royal Society University Research Fellowship (URF\R1\180075) and BBSRC grant BB/T015349/1 to S.S. and by the BBSRC Institute Strategic Program grant BB/J004561/1 to the John Innes Centre.

## Author contributions

B.C., S.S. and M.P. conceived the project. B.C. conducted cryoFIB milling and cryoET; B.C. optimized the sample preparation, collected and processed the cryoEM data, reconstructed the cryoEM map, built and refined the structural models, performed correlative cryo-light and electron microscopy, determined sporulation efficiency and actinorhodin production; B.C and S.S. conducted fluorescent light microscopy; S.S. generated *Streptomyces* strains; J.W.S. and S.S. performed automated Western blot analyses; B.C., J.W.S, S.S. and M.P. wrote the manuscript.

## METHODS

### Bacterial strains, plasmids, and oligonucleotides

Bacterial strains, plasmids, and oligonucleotides can be found in Supplementary Tables 3-4. *E. coli* strains were cultured in LB, SOB, or DNA medium. *E. coli* cloning strains TOP10 and DH5α were used to propagate plasmids and cosmids. *E. coli* strain BW25113/pIJ790 was used for recombineering cosmids^38^. For interspecies conjugation, plasmids were transformed into *E. coli* ET12567/pUZ8002. Where necessary, media was supplemented with antibiotics at the following concentrations: 100 µg/ml carbenecillin, 50 µg/ml apramycin, 50 µg/ml kanamycin, 50 µg/ml hygromycin.

*Streptomyces coelicolor* and *Streptomyces venezuelae* strains were cultivated in LB, MYM, TSB, TSB-YEME, or R2YE liquid medium at 30 °C in baffled flasks or flasks with springs, at 250 rpm or grown on LB, MYM, SFM, R2YE medium solidified with 1.5% (w/v) Difco agar^39^.Where necessary, media was supplemented with antibiotics at the following concentrations: 25 µg/ml apramycin, 5 µg/ml kanamycin, 25 µg/ml hygromycin, 12.5-25 µg/ml nalidix acid.

### Generation of *Streptomyce*s mutant strains

The λ RED homologous recombination system was used to isolate gene replacement mutations using PCR-directed mutagenesis (ReDirect) of the *S. coelicolor* cosmid StD-49 and the *S. venezuelae* cosmid Pl1-F14, containing the CIS gene cluster^38, 40^. Genes encoding the sheath (*sco4253*, *vnz_28920*) or the whole CIS-sheath operon (*sco4253-SCO4251*, *vnz_28920-28910*) were replaced with the *aac3(IV)-oriT* resistance cassette from pIJ773. Mutagenized cosmids (pSS480, pSS481, pSS489, pSS490) were transformed and subsequently conjugated from *E. coli* ET12567/pUZ8002 to wild-type *S. coelicolor* or *S. venezuelae*. Exconjugants that had successfully undergone double-homologous recombination were identified by screening for apramycin-resistance and kanamycin sensitivity. Deletion of the respective CIS mutant genotypes were subsequently verified by PCR.

### Phylogenetic analysis

The phylogenetic analysis of the different contractile injection systems (from eCIS, T6SS, phage and CIS from *Streptomyces*) were examined using the putative sheath proteins. Alignment and generation of the phylogenetic tree was performed as previously reported^16, 19^. First, the amino acid sequences from 16 sheath proteins were aligned by the MUSCLE online tool^41, 42^. Standard parameters were applied for multiple sequence alignment. Then, MEGAX program^43^ was used to reconstruct phylogenetic trees using the Maximum Likelihood (ML) method and bootstrap values (1000 resamples) were applied to assess the robustness of the tree.

### Sheath preparation of CIS from *Streptomyces* for negative-stain EM and mass spectrometry

*S. venezuelae* was cultivated either in 30 mL LB or MYM liquid medium for 14 hours and of *S. coelicolor* strains were grown in 30 ml TSB, TSB-YEME or R2YE liquid medium for 48 hours, respectively. *Streptomyces* cultures were pelleted by centrifugation (7000xg, 10 min, 4 °C), resuspended in 5 ml lysis buffer (150 mM NaCl, 50 mM Tris-HCl, 0.5×CellLytic B (Sigma-Aldrich), 1 % Triton X-100, 200 µg/ml lysozyme, 50 μg/ml DNAse I, pH 7.4), and incubated for 1 hour at 37°C. Cell debris was removed by centrifugation (15000×g, 15 min, 4 °C) and cleared lysates were subjected to ultra-centrifugation (150000×g, 1 h, 4 °C). Pellets were resuspended in 150 µl resuspension buffer (150 mM NaCl, 50 mM Tris-HCl, supplemented with protease inhibitor cocktail (Roche), pH 7.4). Proteins in the CIS preparation were subjected to negative stain EM imaging^44^ and mass spectrometry at the Functional Genomics Center Zürich.

### Negative stain electron microscopy

4 µl of purified sheath particles were adsorbed to glow-discharged, carbon-coated copper grids (Electron Microscopy Sciences) for 60 s, washed twice with milli-Q water and stained with 2 % phosphotungstic acid for 45 s. The grids were imaged at room temperature using a Thermo Fisher Scientific Morgagni transmission electron microscope (TEM) operated at 80 kV.

### Mass spectrometry analysis

To confirm the presence of predicted CIS components from *Streptomyces*, isolated sheath particles were subjected to liquid chromatography–mass spectrometry analysis (LC–MS/MS). First, the samples were digested with 5 µl of trypsin (100 ng/µl in 10 mM HCl) and microwaved for 30 min at 60 °C. The samples were then dried, dissolved in 20 µl ddH_2_0 with 0.1% formic acid, diluted in 1:10 and transferred to autosampler vials for liquid chromatography with tandem mass spectrometry analysis. A total of 1 µl was injected on a nanoAcquity UPLC coupled to a Q-Exactive mass spectrometer (ThermoFisher). Database searches were performed by using the Mascot swissprot and tremble_streptomycetes search program. For search results, stringent settings have been applied in Scaffold (1% protein false discovery rate, a minimum of two peptides per protein, 0.1% peptide false discovery rate). The results were visualized by Scaffold software (Proteome Software Inc., Version 4.11.1).

### Automated Western blot analysis

Automated Western blot analysis (WES) of liquid grown Streptomyces strains was essentially performed as described previously^45^. Cell pellets were resuspended in 0.4 ml of sonication buffer (20 mM Tris pH 8.0, 5 mM EDTA, 1x EDTA-free protease inhibitors [Sigma Aldrich]) and subjected to sonication at 4.5-micron amplitude for 7 cycles of 15 seconds on/15 seconds off. Samples were centrifuged at 14,000 RPM for 15 minutes at 4°C. The supernatants were removed and subjected to a Bradford Assay (Biorad). Equivalent total protein concentrations (0.2 mg/ml) were assayed using the automated Western blotting machine WES (ProteinSimple, San Jose, CA) according to the manufacturer’s guidelines. For the detection of Cis1a and Cis2 protein, antibodies for α-Cis1a (GenScript) and α-Cis2 (GenScript) were used at a concentration of 1:200. For detection of WhiA 0.5 μg of total protein and anti-WhiA (Polyclonal, Cambridge Research Biochemicals) at 1:100 dilution was used^46^.

For the detection of Cis1a and Cis2 in culture supernatants, *S. coelicolor* WT, SS387 and SS395 were grown in duplicate in TSB medium for 48 h. Cultures were pelleted and 20 ml supernatant obtained from each culture were concentrated to approximately 1 ml using Amicon Ultra-15, 10K spin column (Millipore). Total protein samples were further processed as described above. In parallel, an aliquot of each sample was loaded onto a 12 % Teo-Tricine/SDS precast protein gel (Expedian) to demonstrate the presence of proteins in the culture supernatants. SDS-gels were stained with InstantBlue (Sigma-Aldrich) and scanned.

For the automated Western blot analysis of surface-grown *S. coelicolor* samples from R2YE plates, mycelium was scraped of sterile cellophane discs that had been placed on top of solid R2YE medium. Mycelia were removed at the described time points and washed with 1X PBS. The supernatant was discarded and the pellet frozen. Pellets were treated and WES ran as above. All virtual Western blots were generated using the Compass software for simple western (Version 6.0.0). Data of protein abundance was plotted using GraphPad Prism (Version 9.3.1).

For WES analyses of Cis1a and Cis2 abundance following nisin stress *S. coelicolor* WT. cultures were grown in TSB medium at 30 °C for 48 hours, after which they were split and normalized to the same optical density. To one culture replicate, nisin was added to a final concentration of 1 µg/ml and to the other, the diluent (0.05% Acetic acid) was added in equal volume. After which 2 ml aliquots were removed from each sample and pelleted at 13,000 RPM. Pellets were treated as above but were additionally probed with an α-WhiA antibody at 1:100 concentration. The band intensities for Cis1a and Cis2 were normalized against the band intensity of WhiA and plotted in GraphPad Prism (Version 9.3.1) with the standard deviation.

### Fluorescence light microscopy and image analysis

For imaging protein localization and fluorescent promoter reporter fusion in *S. coelicolor*, a Zeiss Axio Observer Z.1 inverted epifluorescence microscope fitted with a sCMOS camera (Hamamatsu Orca FLASH 4), a Zeiss Colibri 7LED light source, a Hamamatsu Orca Flash 4.0v3 sCMOS camera, and a temperature-controlled incubation chamber was used. Images were acquired using a Zeiss Alpha Plan-Apo 100x/1.46 Oil DIC M27 objective with a YFP excitation/emission bandwidths of 489–512 nm/520–550 nm. Still images and time-lapse images series were collected using Zen Blue (Zeiss) and analyzed using Fiji^47^.

To monitor the activity of the fluorescent sheath promoter fusion in *S. coelicolor*, spores of strain SS484 were spotted onto solid R2YE medium and grown alongside a microscopic coverslips that had been inserted into the agar at an approximately 45 ° angle. Plates were incubated at 30 °C for up to 4 days. At the indicated time points, glass coverslips with attached hyphae were removed and mounted onto slides affixed with 1 % agar pads and imaged.

For time-lapse imaging of *S. coelicolor* expressing a fluorescently labelled sheath protein (SS389), cells were first grown in TSB-YEME for 40 h and a 2 μl sample of the culture was immobilized on a 1 % agarose pad prepared with filtered culture medium and using a Gene Frame (Thermo Scientific). Experiments were performed at 30 °C and growing hyphae were imaged every 5 min. Image collection and analysis was performed using Zen Blue (Zeiss) and Fiji, respectively^47^.

### Plunge freezing of *Streptomyces* hyphae

For cryo-electron tomography (cryoET), *Streptomyces* cells were mixed with 10 nm Protein A conjugated colloidal gold particles (1:10 v/v, Cytodiagnostics) and 4 µl of the mixture was applied to a glow-discharged holey-carbon copper EM grid (R2/1 or R2/2, Quantifoil). The grid was automatically blotted from the backside for 4-6 s in a Mark IV Vitrobot by using a Teflon sheet on the front pad, and plunge-frozen in a liquid ethane-propane mixture (37%/63%) cooled by a liquid nitrogen bath.

For single particle cryoEM (SPA), the *S. coelicolor* CIS particles (from WT CIS and non-contractile CIS), collected after sheath preparation, were vitrified using a Vitrobot Mark IV (Thermo Fisher Scientific). 4 µl of samples were applied on glow-discharged 200 mesh Quantifoil Gold grids (R 2/2). Grids were blotted for 5 s and plunged into liquid ethane-propane mix (37 %/63 %). Frozen grids were stored in liquid nitrogen until loaded onto the microscope.

### Cryo-focused ion beam milling

A standard protocol was used to perform cryo-focused ion beam milling (CryoFIB milling) on *S. venezuelae*^48^. Plunge-frozen grids were clipped into cryoFIB-autoloader grids (Thermo Fisher Scientific), then transferred into a liquid nitrogen bath of a loading station (Leica Microsystems) and mounted into a 40 ° pre-tilted SEM grid holder (Leica Microsystems). The holder was transferred with a VCT100 cryo-transfer system (Leica Microsystems) into a Helios NanoLab600i dual beam FIB/scanning electron microscope (SEM, Thermo Fisher Scientific). Grids were coated with platinum precursor gas for 6 s and checked with SEM at 3-5 kV (80 pA) to evaluate grid quality and identify targets. Lamella were milled in multiple steps using the focused gallium ion beam (43 nA to 24 pA) until a thickness ∼250 nm was achieved. The holder was returned to the loading station using the VCT100 transfer system. Unloaded grids were stored in liquid nitrogen prior to cryoET imaging.

### Cryo-electron tomography

Intact or cryoFIB-milled *Streptomyces* cells were imaged by cryoET^49^. Images were recorded on Titan Krios 300 kV microscopes (Thermo Fisher Scientific) equipped with a Quantum LS imaging filter operated at a 20 eV slit width and with K2 or K3 Summit direct electron detectors (Gatan). Tilt series were collected using a bidirectional tilt-scheme from −60 to +60 ° in 2 ° increments. Total dose was 130-150 e^-^/Å^2^ and defocus was kept at −8 µm. Tilt series were acquired using SerialEM^50^, drift-corrected using alignframes, reconstructed and segmented using IMOD program suite^51^. To enhance contrast, tomograms were deconvolved with a Wiener-like filter^52^.

### SPA data collection and image processing

CryoEM datasets of *S. coelicolor* contracted sheath and extended sheath-tube module were collected as movie stacks using the SerialEM program on Titan Krios EM operating at 300 kV and equipped with an energy filter and a K2 Summit camera. The movie frames of each collected stack were aligned and summed up into one single micrograph with dose weighting at the binning factor of 2 using MotionCor2 . The CTF parameter of the micrographs were estimated using Gctf. Pixel size at specimen level was 1.4 Å and target defocus ranged from 1.5 µm to 3.5 µm. Each stack contains 50 frames, and the accumulated electron dose rate was ∼60 e^-^/Å^2^.

The image processing of contracted sheath and extended sheath-tube from *S. coelicolor* was performed as previously reported^19^. The particles were picked manually using Relion 3.0^54^. The particle extraction was performed in “Extract helical segments” mode to extract helical segments. The structural determination of the contracted sheath and the extended sheath-tube module was performed using helical reconstruction in Relion 3.0^55^.

For the contracted sheath, the final 3.6 Å resolution structure of contracted sheath was obtained from 4,838 particles applied with 6-fold symmetry and helical parameters (rise = 17.22 Å, twist = 26.58 °) (Extended Data Fig. 2a).

For the extended sheath-tube module, the final 3.9 Å resolution structure of the extended sheath-tube module was determined from 18,822 particles calculated with 6-fold symmetry and helical parameters (rise = 38.50 Å, twist = 23.10 °) (Extended Data Fig. 3e).

The resolutions of relative reconstruction maps were estimated based on the gold-standard Fourier Shell Correlation (FSC) = 0.143 criteria^56^. The local resolution estimations of individual maps were performed using the local resolution module in Relion 3.0 and examined using UCSF Chimera^57^ (Extended Data Fig. 2b and Extended Data Fig. 3f).

### Structure modeling

Proteins were built *de novo* using COOT^58^. Models were iteratively refined using RosettaCM^59^ and real-space refinement implemented in PHENIX^60^. Sheath protein could only be partially modeled and in some cases side chains were not assigned. Final model validation was done using MolProbity^60^ and correlation between models and the corresponding maps were estimated using mtriage^60^.

All visualizations were done using PyMOL, UCSF Chimera^57^ or ChimeraX^61^.

### Correlative cryo-light and electron microscopy

For correlative cryo-light and electron microscopy, frozen grids containing *S. coelicolor* WT were transferred to CMS196V3 Linkam cryo-stage and imaged using a 100x numerical aperture 0.74 objective on a LSM900 Airyscan 2 Zeiss microscope driven by ZEN Blue software (Version 3.5). Fluorescence images of areas of interest were manually correlated with the corresponding TEM square montage using SerialEM^50, 62^.

### Fluorescence-based cell viability assay

To express sfGFP constitutively in *Streptomyces* strains, the coding sequence for sfGFP was introduced downstream of the constitutive promoter *ermE*\* on an integrating plasmid vector (pIJ10257). The plasmid was introduced by conjugation to *S. coelicolor* strains (WT, ΔCIS and CIS-N5). These strains were inoculated into 30 ml of TSB liquid culture and incubated at 30 °C with shaking at 250 rpm in baffled flasks for 48 h. Where appropriate, nisin and CCCP (or 0.002% DMSO) were added to a final concentration of 1 µg/ml and 10 µM, respectively. Cultures were incubated for a further 90 min. For UV exposure, 10 ml of the *S. coelicolor* cultures were transferred into a petri dish and treated with Sankyo Denki Germicidal 68 T5 UV-C lamps for 10 mins in a Herolab UV DNA crosslinker CL-1. Then, 1 ml aliquots were centrifuged for 5 min at 13,000 rpm, washed twice with PBS, and resuspended in 1 ml of PBS with 5 µg/ml FM5-95 membrane stain. The cell suspension and membrane stain were mixed by vortexing and kept in the dark at room temperature for 10 min. The suspension was then centrifuged for 5 min at 13,000 rpm, washed twice with PBS, and resuspended in 50 µl of PBS. 10 µl of samples were immobilized on 1% agar pads and imaged on the Thunder imager 3D cell culture microscope at room temperature. First, tile scan images were acquired on the Las X Navigator plug-in of Leica Application Suite X (LasX) software (Version 3.7.4.23463), and 100 targets were picked manually. Then z-stack images with HC PL APO 100x objective were acquired at an excitation of 475 nm and 555 nm under GFP (green) and TRX (red) filters respectively. Images were processed using LasX software to apply thunder processing and maximum projection, FIJI to create segmentation and quantify the live (sfGFP)/total cells (FM5-95) area ratio^47^ and statistical analysis was performed on GraphPad Prism 9 (Version 9.3.1).

### Cover glass impression of *Streptomyces* spore chains

Spore titers of relevant strains were determined by standard techniques. 10^7^ CFU of *S. coelicolor* strains (WT, SS387, SS393 and SS395) were spread onto R2YE agar plates and grown at 30 °C. Sterile glass cover slips were gently applied to the top surface of each bacterial lawn after 48 h, 72 h and 96 h post inoculation. Cover slips were then mounted onto glass microscope slides and imaged using a 40x objective on a Leica Thunder Imager 3D Cell Culture. Images were processed using FIJI^47^.

### Actinorhodin production assay

*S. coelicolor* strains (WT, SS387, SS393 and SS395) were inoculated into 30 ml R2YE liquid media at a final concentration of 1.5 x 10^6^ CFU/ml. Cultures were grown in baffled flasks at 30 °C overnight. Cultures were standardized to an OD_450_ of 0.5 and inoculated in 30 ml of fresh R2YE liquid medium. For visual comparison of pigment production, images of the growing culture were taken between t = 0 and t = 72 h (as indicated in Extended Data Fig. 9c). For quantification of total actinorhodin production, 480 µl of samples were collected at the same time points where images were taken. 120 µl of 5M KOH was added, samples were vortexed and centrifuged at 5000 x g for 5 min. The weight of each tube was recorded. A Synergy 2 plate reader (Biotek) was used then to measure the absorbance of the supernatant at 640 nm. The absorbance was normalized by weight of the wet pellet.

## Data availability

Representative reconstructed tomograms (EMD-XXXXX, EMD-XXXXX, EMD-XXXXX, EMD-XXXXX, EMD-XXXXX, EMD-XXXXX and EMD-XXXXX) and SPA cryoEM maps (EMD-XXXXX and EMD-XXXXX) have been deposited in the Electron Microscopy Data Bank. Atomic models (PDB: XXXX and PDB: XXXX) have been deposited in the Protein Data Bank. All other data are available from the authors upon reasonable request.

---

**Extended Data Figure 1:**
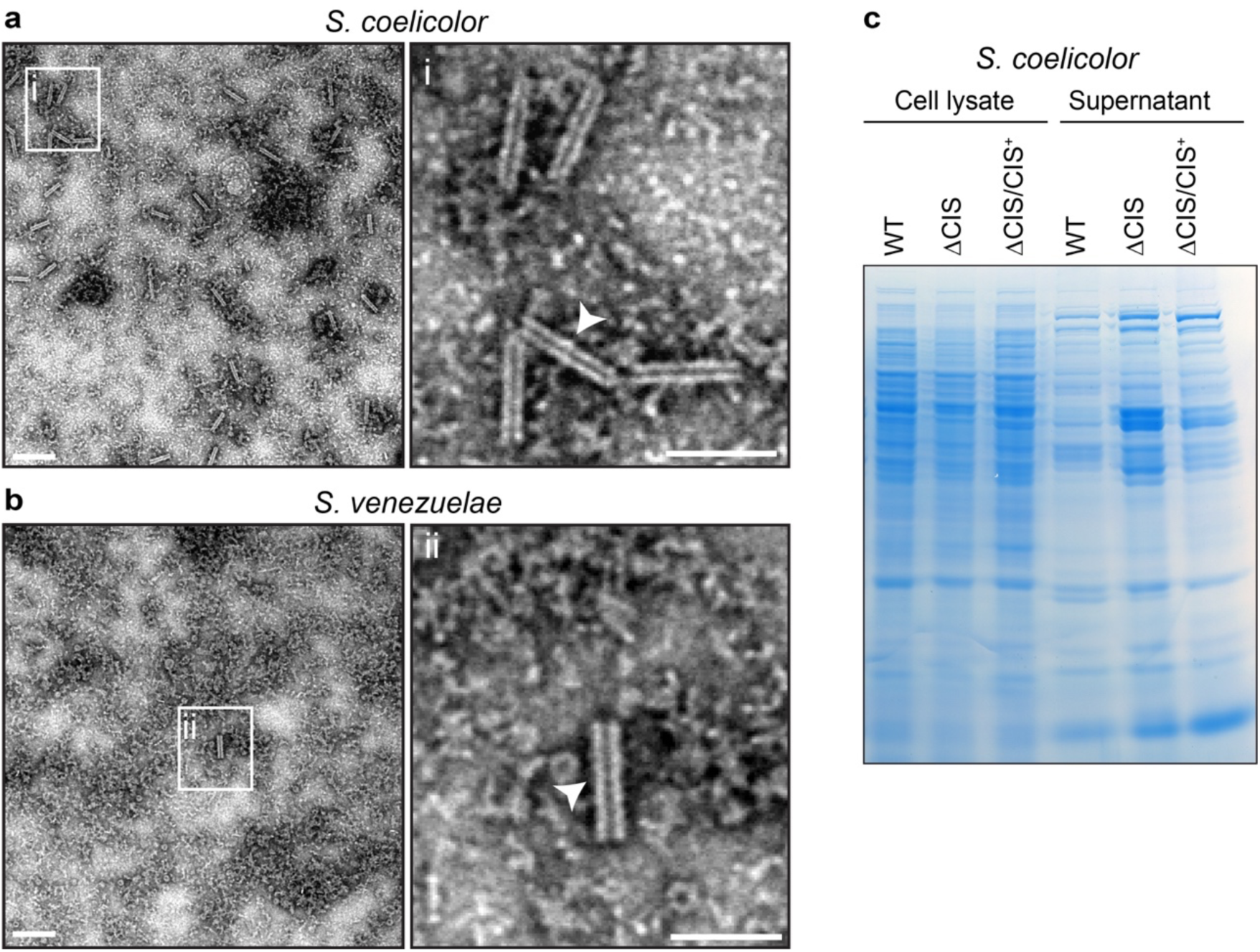
EM and SDS-PAGE analyses of *Streptomyces* supernatant and lysate. **a/b.** Representative negative-stain electron micrographs of crude sheath preparations from WT *S. coelicolor* and *S. venezuelae.* Under the conditions used, the majority of isolated CIS from *Streptomyces* was contracted (insets i/ii). Bars, 80 nm. **c.** Control SDS-PAGE stained with Coomassie-blue showing the presence of protein in concentrated culture supernatants that were used for the detection of Cis1a/2 by automated Western blot analysis in Figure 1e. Samples were obtained from WT *S. coelicolor*, the 11CIS mutant and the complemented mutant 11CIS/CIS^+^. Loaded were 10 μg of protein per sample.

**Extended Data Figure 2:**
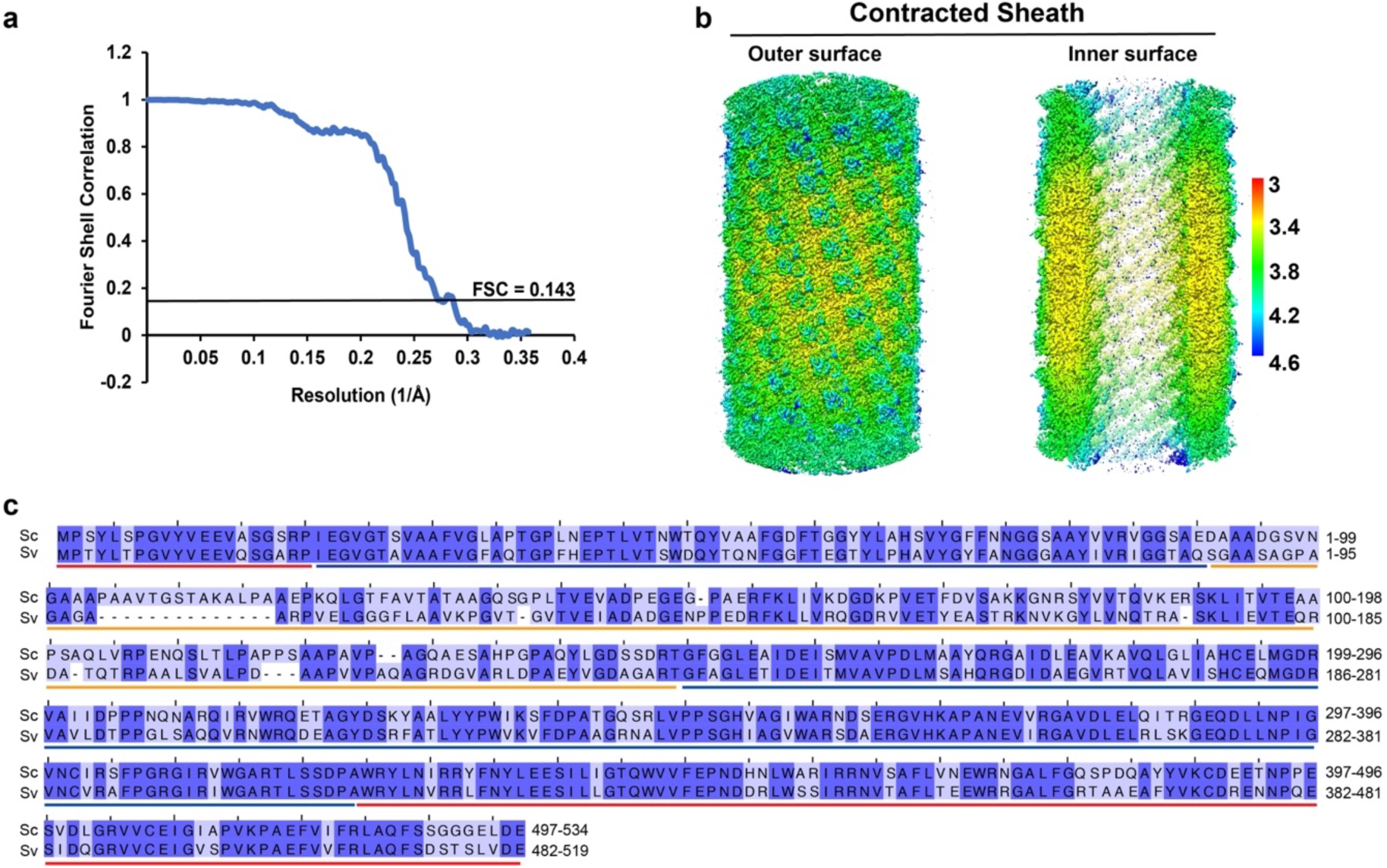
Structure and sequence analysis of *Streptomyces* contracted sheath Cis2. **a/b.** Gold-standard Fourier shell correlation (FSC) curve (a) and local resolution maps (b) of the contracted sheath (Cis2) structures from *S. coelicolor*. **c.** Protein sequence alignment showing the high sequence conservation between Cis2 proteins from *S. coelicolor* (Sc) and *S. venezuelae* (Sv). Colors indicate level of sequence similarity (light blue, similar; dark blue, identical). Positions of domain 1 (red), domain 2 (blue) and domain 3 (orange) are indicated.

**Extended Data Figure 3:**
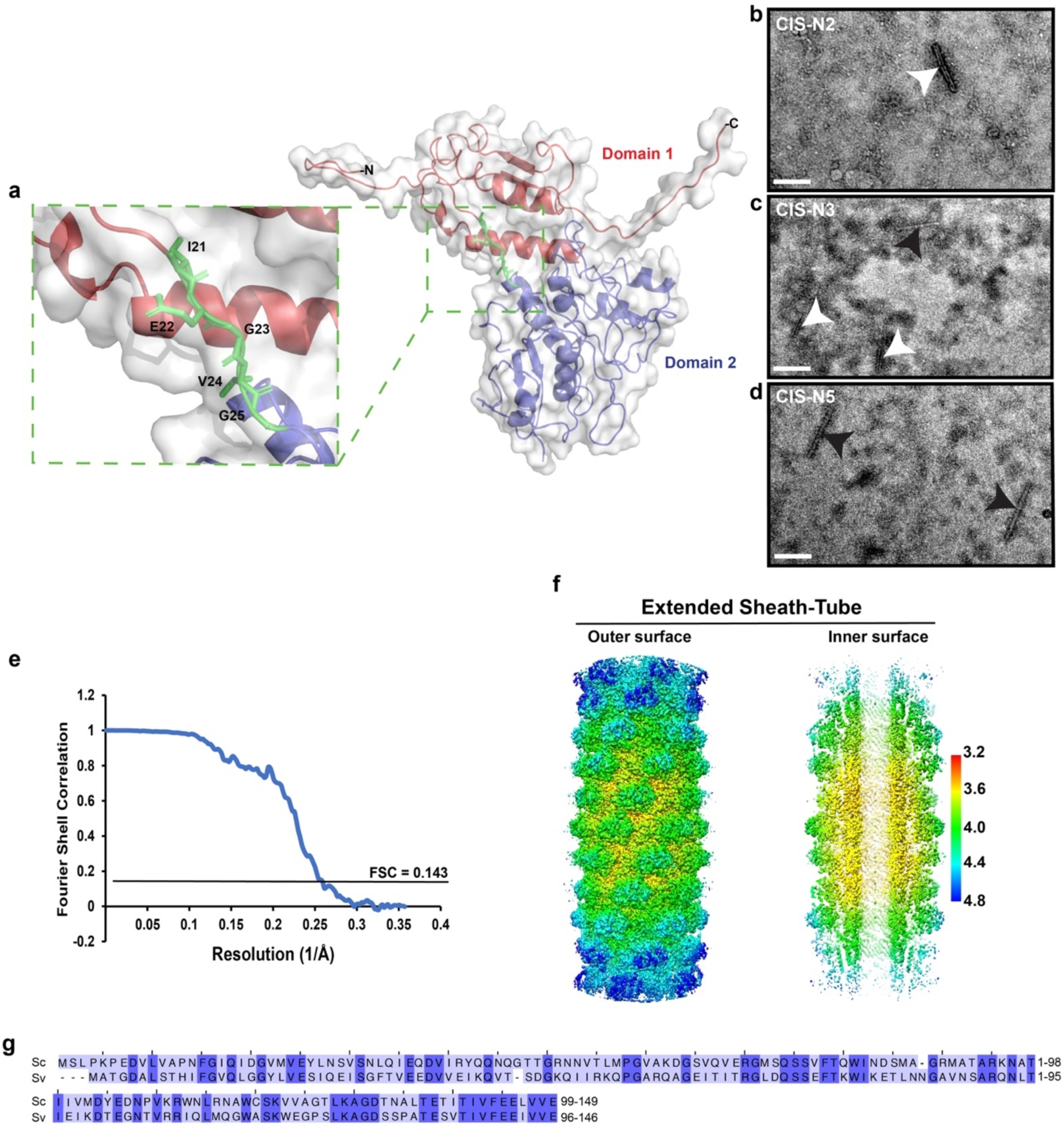
Engineering and structure of a non-contractile CIS^Sc^ mutant. **a.** Surface (grey) and ribbon (colored) representation of the contracted sheath structure from *S. coelicolor.* Shown in the enlarged inset is the WT linker comprising residues I21 to G25 (green). In order to engineer non-contractile CIS^Sc^ mutants, additional residues were inserted after position G25 (IE for CIS*-*N2, IEG for CIS*-*N3, IEGVG for CIS*-*N5). **b-d.** Negative-stain electron micrographs of CIS particles from *S. coelicolor* strains expressing the CIS^Sc^ mutant versions CIS*-*N2, CIS-N3 and CIS-N5. Sheath mutants carrying the CIS-N5 allele showed the highest fraction of extended structures. Arrowheads indicate contracted (white) and extended (black) CIS^Sc^ particles. Bar, 140 nm. **e-f.** Gold-standard Fourier shell correlation (FSC) curve (e) and local resolution maps (f) of the extended *S. coelicolor* CIS^Sc^-N5 sheath-tube module. **g.** Protein sequence alignment showing the high conservation of Cis1a proteins from *S. coelicolor* (Sc) and *S. venezuelae* (Sv). Colors indicate level of sequence similarity (light blue, similar; dark blue, identical).

**Extended Data Figure 4:**
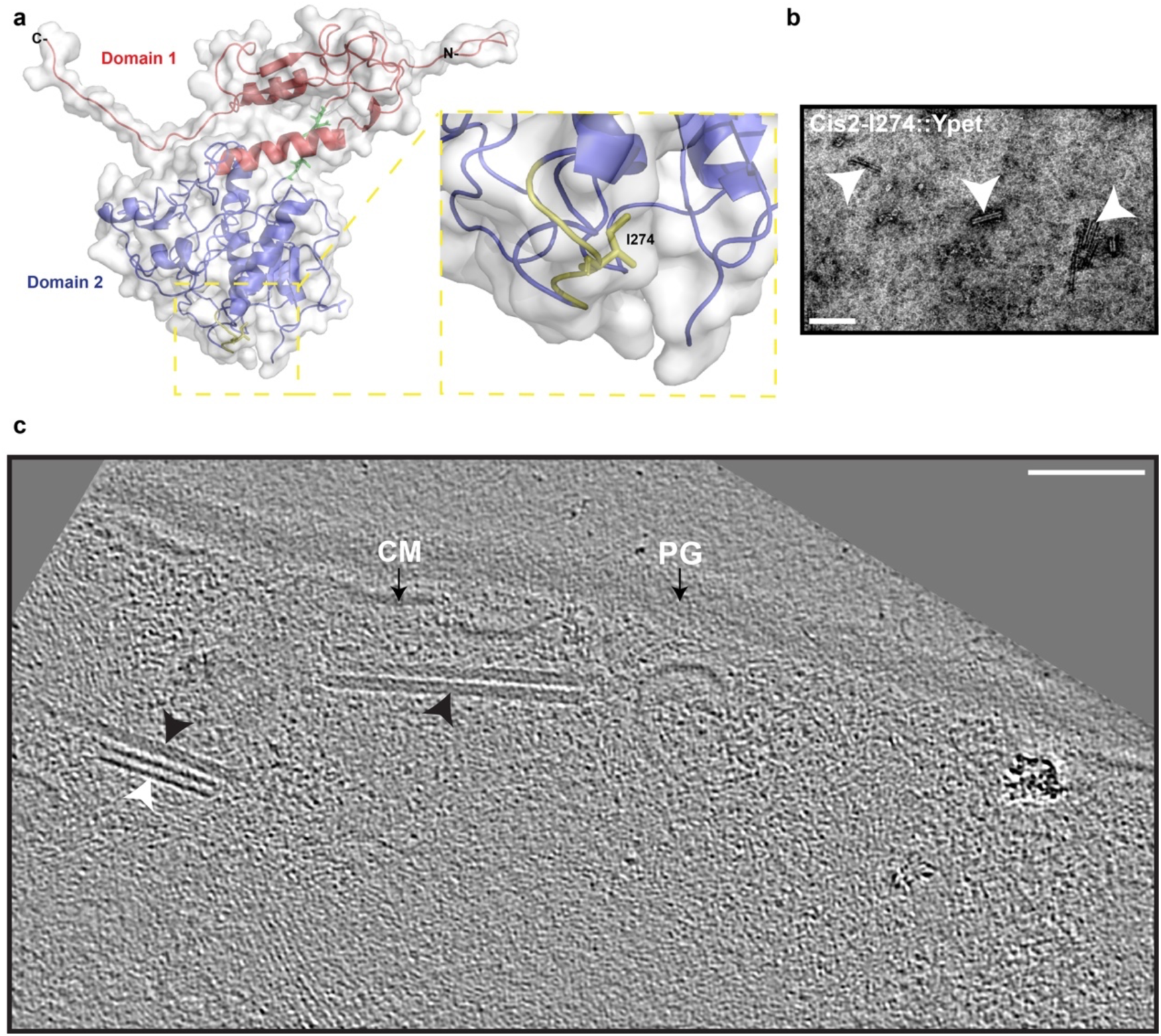
Generation of a functional Cis2-YPet sandwich fusion. **a.** Surface and ribbon diagram of the contracted sheath structure from *S. coelicolor* indicating the insertion site of Ypet at residue I274 to generate a fluorescent sheath-Ypet sandwich fusion (Cis2::I274-Ypet), which was used to complement a *S. coelicolor* 11Cis2 mutant. **b.** Negative-stain electron micrograph of purified CIS particles from *S. coelicolor Δcis2/cis2::I274-ypet^+^* showing contracted CIS^Sc^ assemblies (white arrowheads). Bar, 140 nm. **c.** Representative cryoET slice of *Δcis2/cis2::I274-ypet^+^* hyphae containing a contracted (white arrowhead) and extended (black arrowhead) CIS^Sc^ particles in the cytoplasm. PG, peptidoglycan; CM, cytoplasmic membrane. Bar, 75 nm.

**Extended Data Figure 5:**
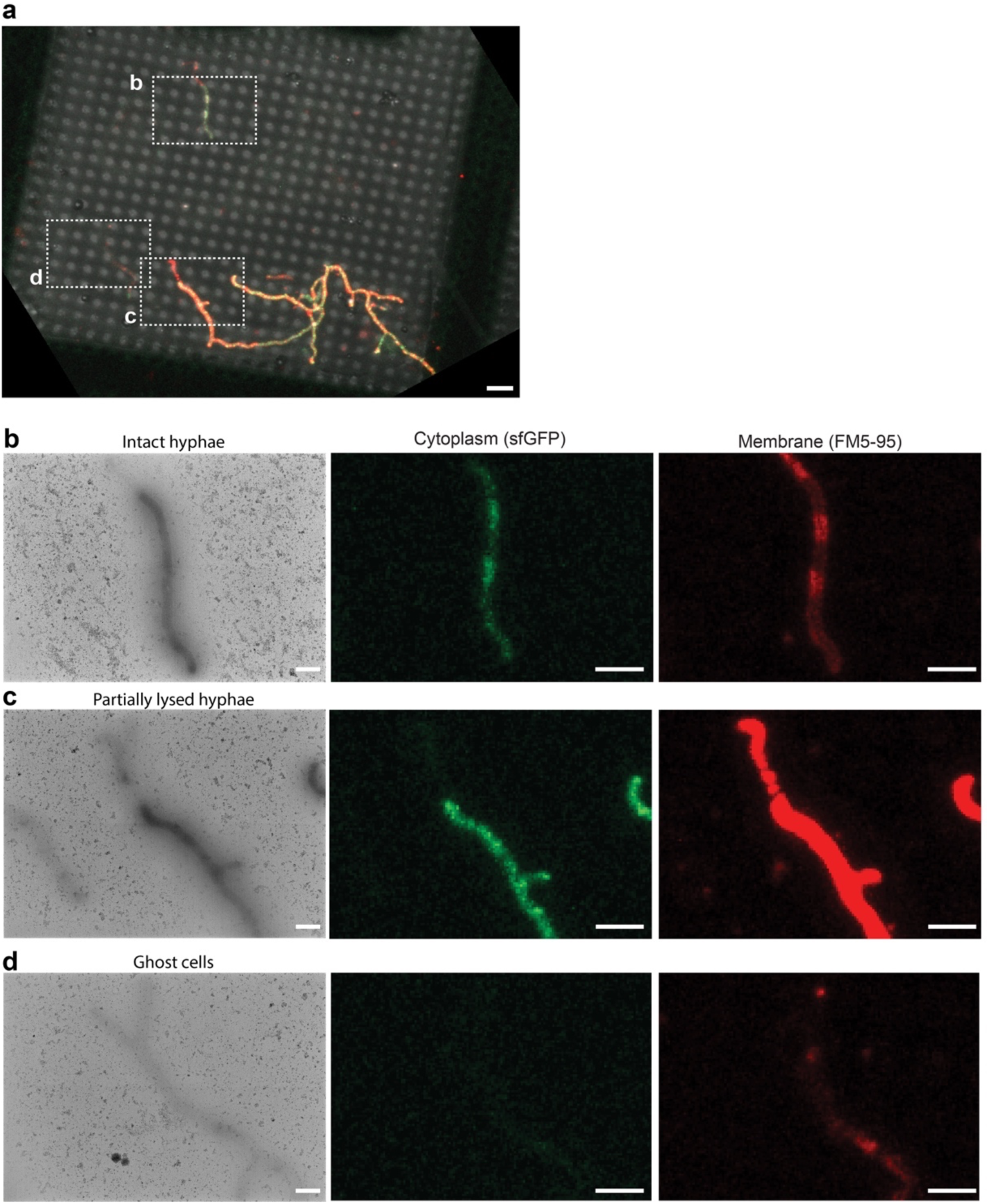
Validation of hyphal membrane integrity using correlative cryo-fLM and cryoEM (Cryo-CLEM). **a.** CryofLM overview image of vegetative hyphae of WT *S. coelicolor* expressing cytoplasmic sfGFP from a constitutive promoter and stained with the membrane dye FM5-95. The membrane staining pattern and sfGFP fluorescence signal were used to identify the classes ‘intact hyphae’, ‘partially lysed hyphae’ and ‘ghost cells’. The boxed areas were further analyzed in (b-d). Bar, 6 µm. **b-d.** Shown are cryoEM 2D projection images (left) and the corresponding cryo-fLM images of examples of ‘intact hyphae’ (b), ‘partially lysed hyphae’ (c) and ‘ghost cells’ (d). Bars, 2 µm.

**Extended Data Figure 6:**
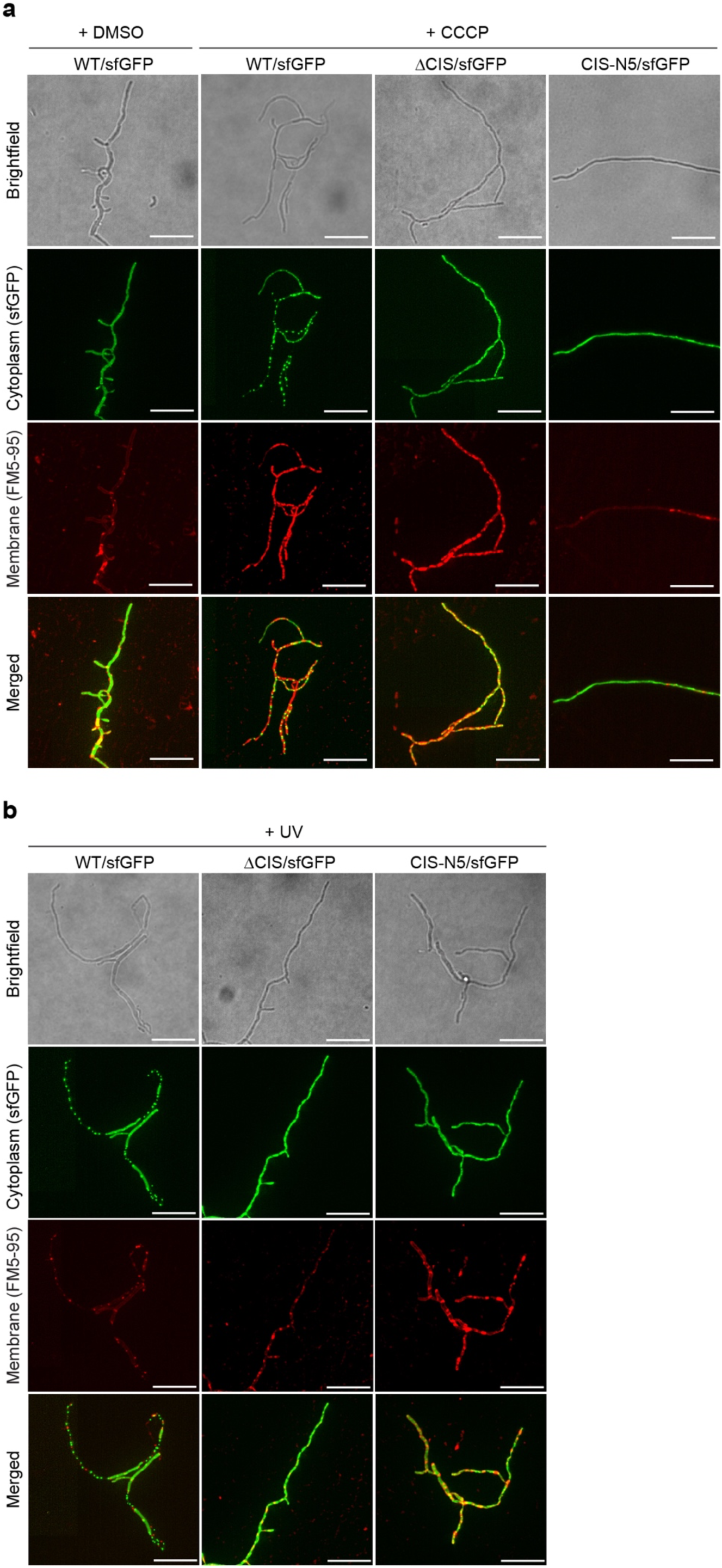
Functional CIS^Sc^ production promotes stress-induced cell death. **a/b.** fLM (shown are representative images) was used to determine the ratio between live cells (cytoplasmic sfGFP) and total cells (membrane dye FM5-95) after growth in the presence of 10 µM CCCP (or 0.002 % DMSO as mock control) (a), or after exposure to UV light (b). *S. coelicolor* WT/sfGFP, ΔCIS/sfGFP and CIS-N5/sfGFP were grown in TSB for 48 h and treated with CCCP or DMSO for 90 min or were exposed to UV light for 10 min. Bars, 10 µm. The quantification for both experiments is shown in Fig. 4 e/f.

**Extended Data Figure 7:**
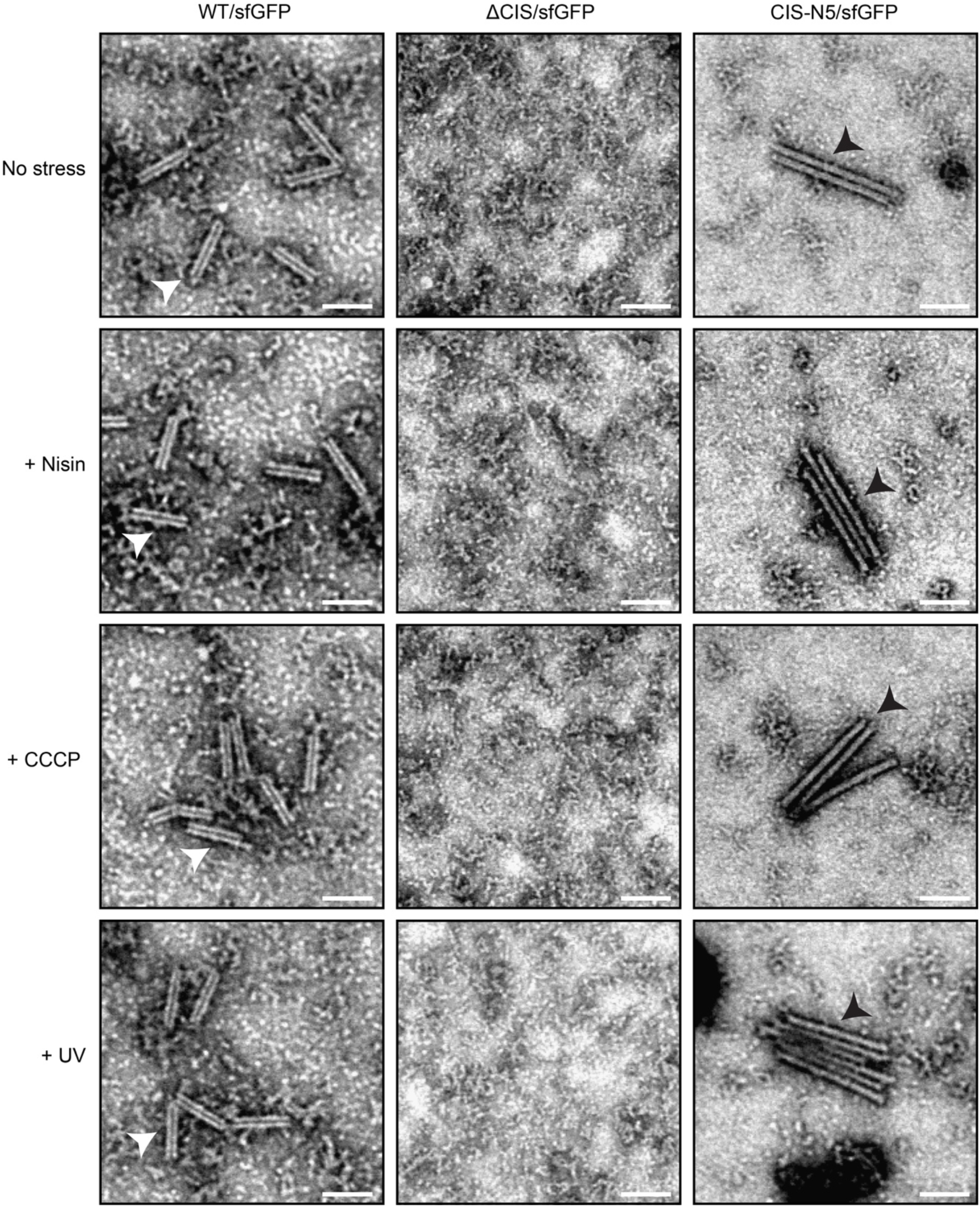
Cell envelope and UV stress do not affect the overall appearance of purified CIS^Sc^ particles. Negative-stain electron micrographs of CIS particles purified from *S. coelicolor* WT/sfGFP, ΔCIS/sfGFP and CIS-N5/sfGFP exposed to no stress, 1µg/ml nisin, 10 µM CCCP and UV treatment. Arrowheads indicate contracted (white) and extended (black) CIS^Sc^. Bars, 90 nm.

**Extended Data Figure 8:**
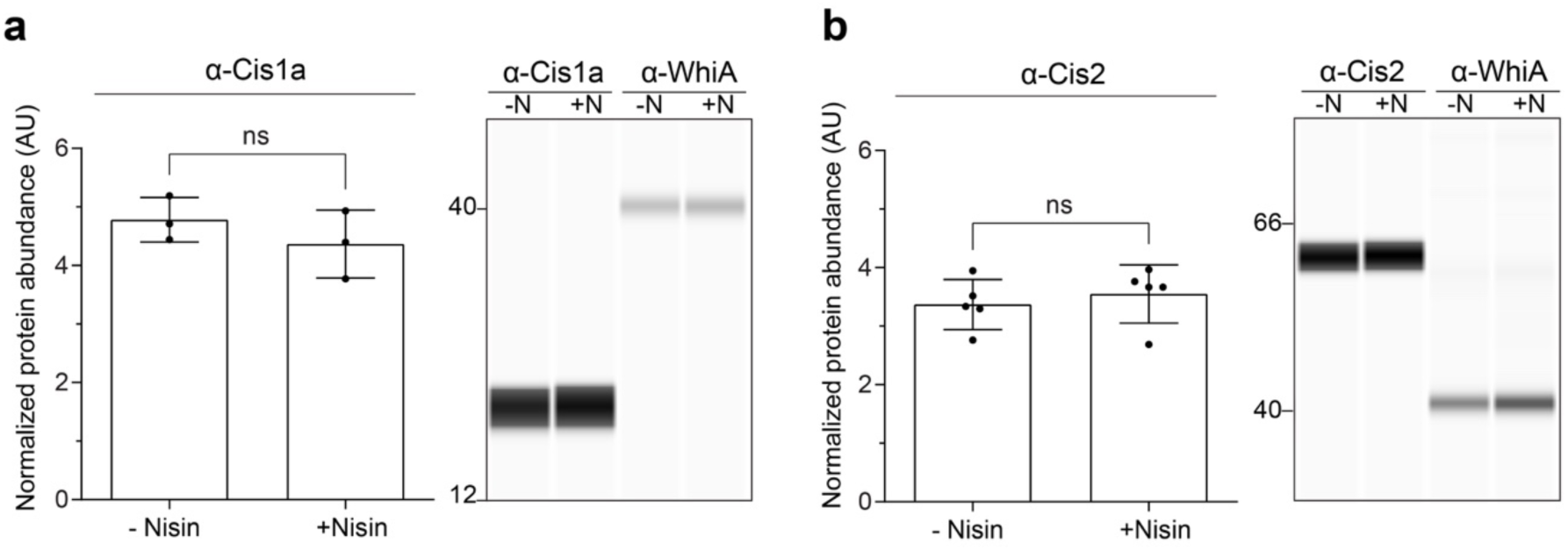
Nisin-stress does not lead to increased CIS^Sc^ production. **a/b.** Quantification (left) and automated Western blot (right) analysis, showing the abundance of Cis1a (a) and Cis2 (b) in WT *S. coelicolor* cell lysates in the presence of nisin (+N) and absence of nisin (-N). Cells were grown in TSB for 48 h, followed by treatment with 1 µg/ml nisin for 90 min. Equal amounts of total protein were subjected to automated Western blot analysis and probed with α-Cis1a, α-Cis2 and α-WhiA polyclonal antibodies. Analysis was performed in biological triplicate experiments. Cis1a/2 protein levels were normalized to WhiA levels. Shown are the mean values and standard error. ns (not significant) and *p*-value was determined using a two-tailed *t*-test.

**Extended Data Figure 9:**
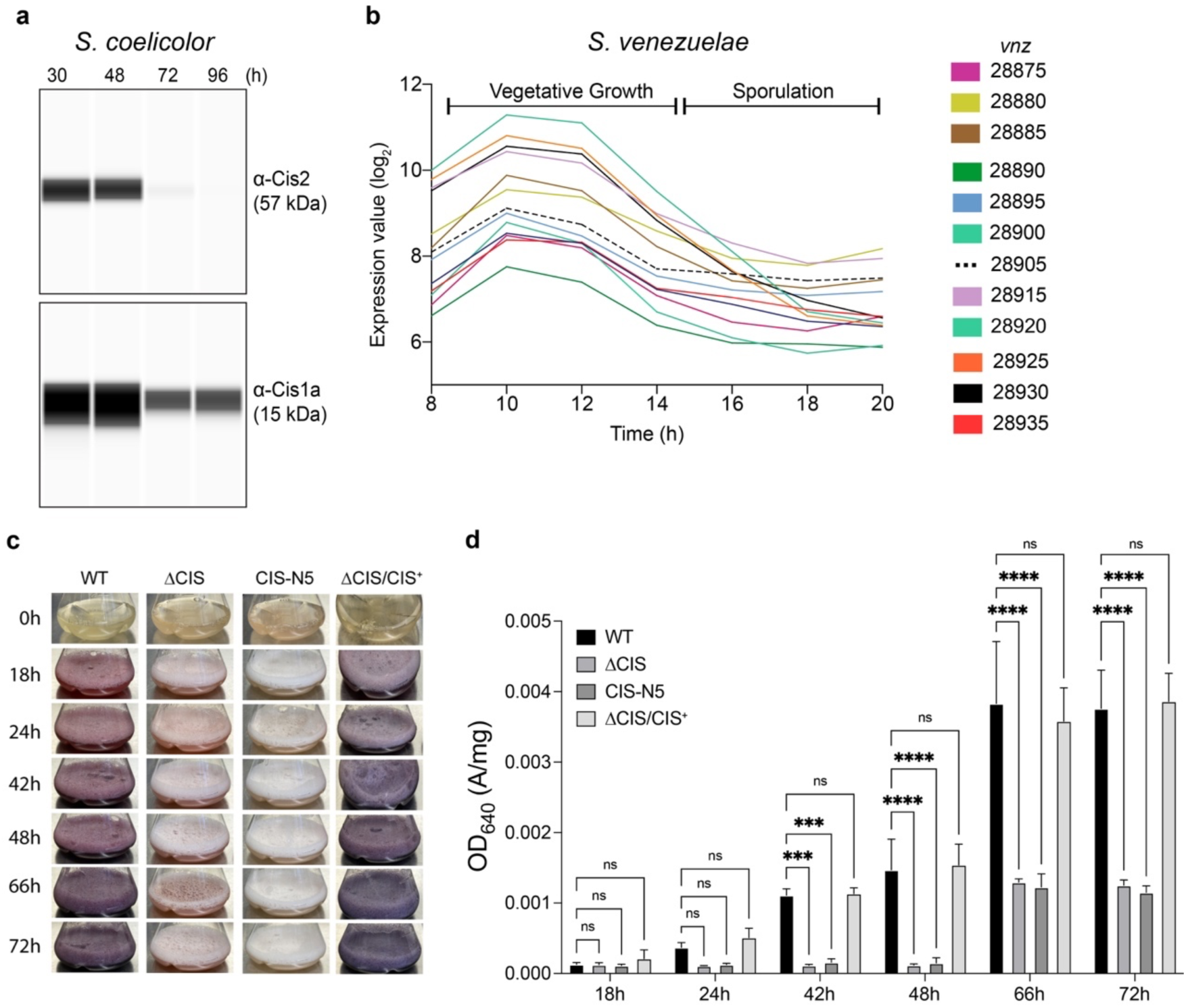
*Streptomyces* CIS proteins are expressed during vegetative growth and impact secondary metabolite production. **a.** Automated Western blot showing the expression of Cis1a (inner tube) and Cis2 (sheath) in hyphae of WT *S. coelicolor* over a time-course of 96 h. *S. coelicolor* was grown on cellophane discs on top of R2YE agar. Automated Western blot analysis was performed in biological duplicate experiments. Equal amounts of protein lysate were loaded and Cis1a and Cis2 were detected using polyclonal α-Cis1a and α-Cis2 antibodies. **b.** Transcription profile of the *S. venezuelae* CIS gene cluster over the entire life cycle^32^. **c.** Comparison of the coloration pattern of *S. coelicolor* WT, the CIS^Sc^ mutant strains ΔCIS, CIS-N5 and the complemented mutant ΔCIS/CIS^+^ in R2YE liquid media. Coloration is indicative of actinorhodin (purple) and undecylprodigiosin (red) production^34^. Note the difference in coloration between WT/complementation mutant as compared to both CIS^Sc^ mutants. Images of each culture flask were taken at the indicated time points. **d.** Quantification of total actinorhodin production of the samples shown in (c). The optical density OD_640_ is an indicator for actinorhodin production^64^. OD_640_ of the culture supernatants was measured and normalized to pellet weight. Note the significant differences that were detected between WT/complementation mutant as compared to both CIS mutants at later time points. Bar plots and error bars represent three biological replicates. p-values (***p < 0.001 and ****p < 0.0001) were calculated using one-way ANOVA and Tukey’s post-test. ns, not significant.

**Supplementary Movie 1.** Time-lapse movie related to Figure 2g/h showing the spatiotemporal localization of fluorescently tagged CIS particles in growing *S. coelicolor* hyphae. Images were acquired every 5 min. Bar, 10 μm.

**Supplementary Table 1.** List of peptides detected by mass spectrometry in samples of purified CIS from WT *S. coelicolor* (ScoWT) and *S. venezuelae* (SvenWT), the corresponding 11Cis2 mutants and the *S. coelicolor* non-contractile CIS^Sc^ mutant CIS-N5. Experiments were performed in biological replicates.

| Protein ID | CIS ID | *S. coelicolor* WT | *S. coelicolor* ΔCis2 | *S. coelicolor* CIS-N5 | *S. venezuelae* WT | *S. venezuelae* ΔCis2 |
| --- | --- | --- | --- | --- | --- | --- |
| Sco4244/Vnz_28875 | Cis11 | - | - | 29% coverage / 10 total unique peptide | - | - |
| Sco4245/Vnz_28880 | Cis9 | - | - | 21% coverage / 2 total unique peptide | - | - |
| Vnz_28885 | Cis10 | - | - | - | - | - |
| Sco4246/Vnz_28890 | Cis8 | - | - | 46% coverage / 18 total unique peptide | - | - |
| Sco4247/Vnz_28895 | Cis7 | - | - | 51% coverage / 8 total unique peptide | - | - |
| Sco4248/Vnz_28900 | Cis5 | - | - | 39% coverage / 4 total unique peptide | - | - |
| Sco4251/Vnz_28915 | - | - | - | - | - | - |
| Sco4252/Vnz_28920 | Cis1 | 27% coverage / 4 total unique peptide | - | 32% coverage / 5 total unique peptide | 17% coverage / 3 total unique peptide | - |
| Sco4253/Vnz_28925 | Cis2 | 48% coverage / 26 total unique peptide | - | 51% coverage / 28 total unique peptide | 38% coverage / 17 total unique peptide | - |
| Sco4254 | - | - | - | 10% coverage / 6 total unique peptide | - | - |
| Sco4259/Vnz_28930 | Cis15 | - | - | - | - | - |
| Sco4260/Vnz_28935 | Cis16 | - | - | 24% coverage / 4 total unique peptide | - | - |

**Supplementary Table 2.**
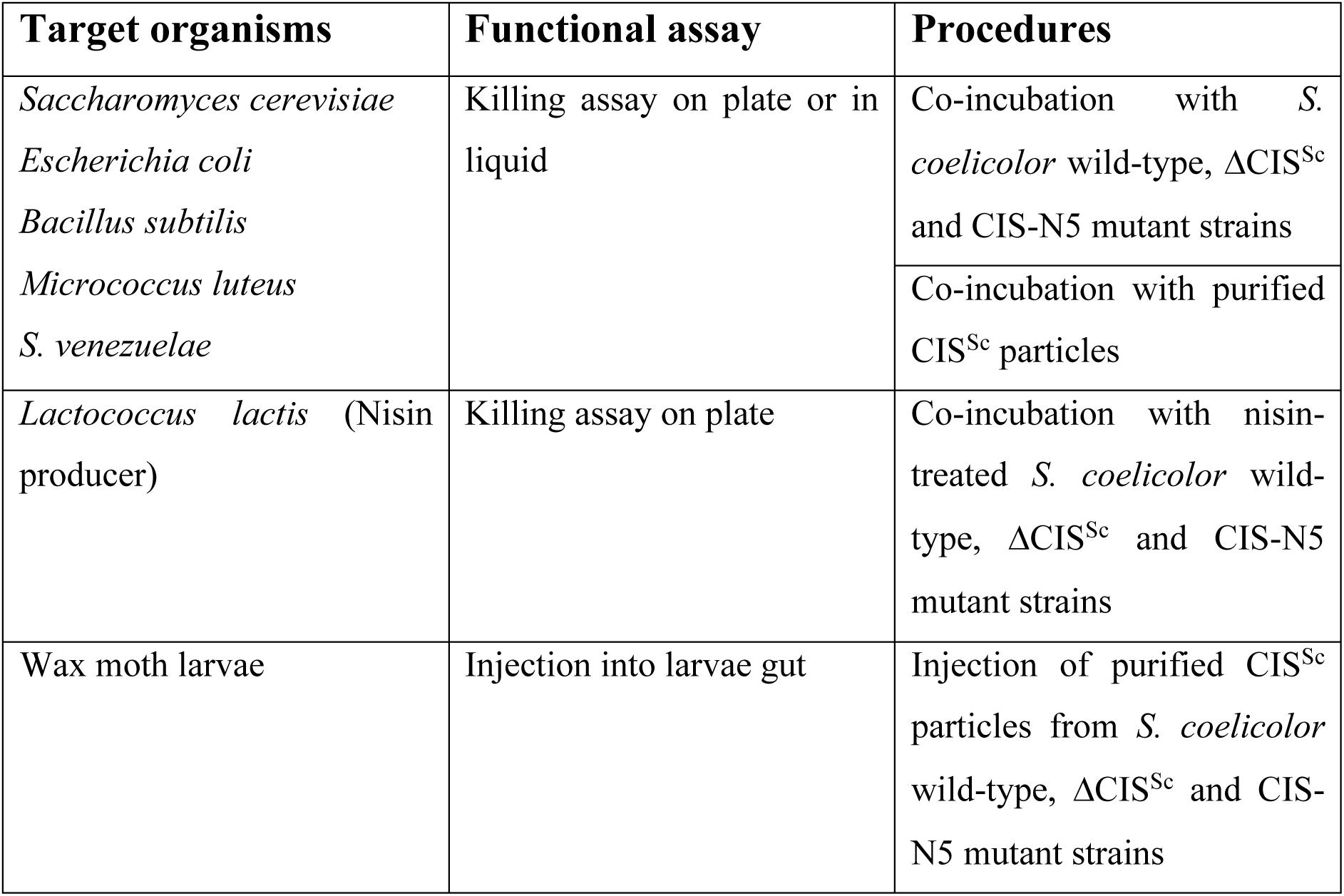
Experimental approaches to study effects of CIS^Sc^ on interspecies competition

| Target organisms | Functional assay | Procedures |
| --- | --- | --- |
| *Saccharomyces cerevisiae*<br>*Escherichia coli*<br>*Bacillus subtilis*<br>*Micrococcus luteus*<br>*S. venezuelae* | Killing assay on plate or in liquid | Co-incubation with *S. coelicolor* wild-type, ΔCIS<sup>Sc</sup> and CIS-N5 mutant strains |
|  |  | Co-incubation with purified CIS<sup>Sc</sup> particles |
| *Lactococcus lactis* (Nisin producer) | Killing assay on plate | Co-incubation with nisin-treated *S. coelicolor* wild-type, ΔCIS<sup>Sc</sup> and CIS-N5 mutant strains |
| Wax moth larvae | Injection into larvae gut | Injection of purified CIS<sup>Sc</sup> particles from *S. coelicolor* wild-type, ΔCIS<sup>Sc</sup> and CIS-N5 mutant strains |

**Supplementary Table 3:**
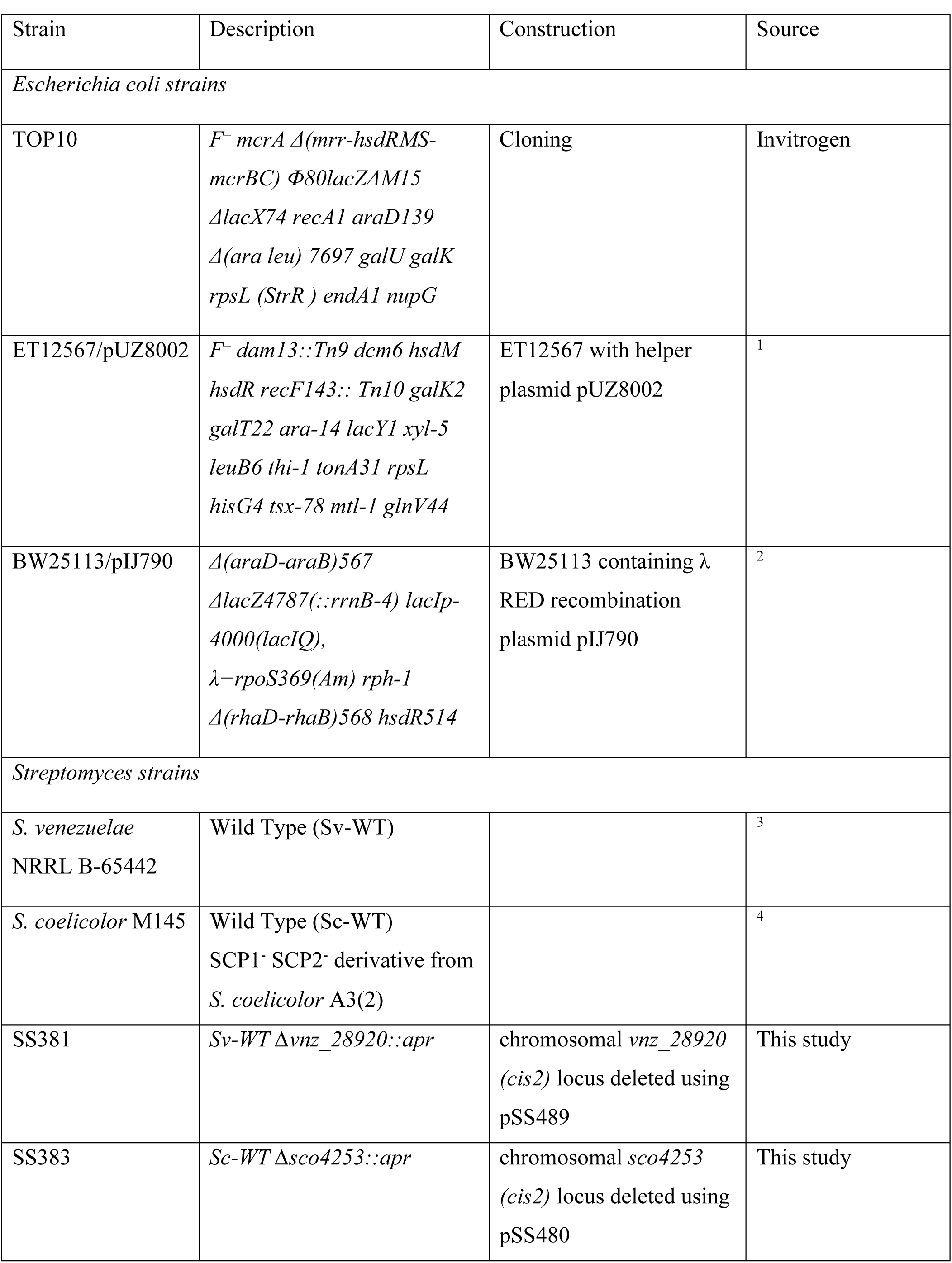
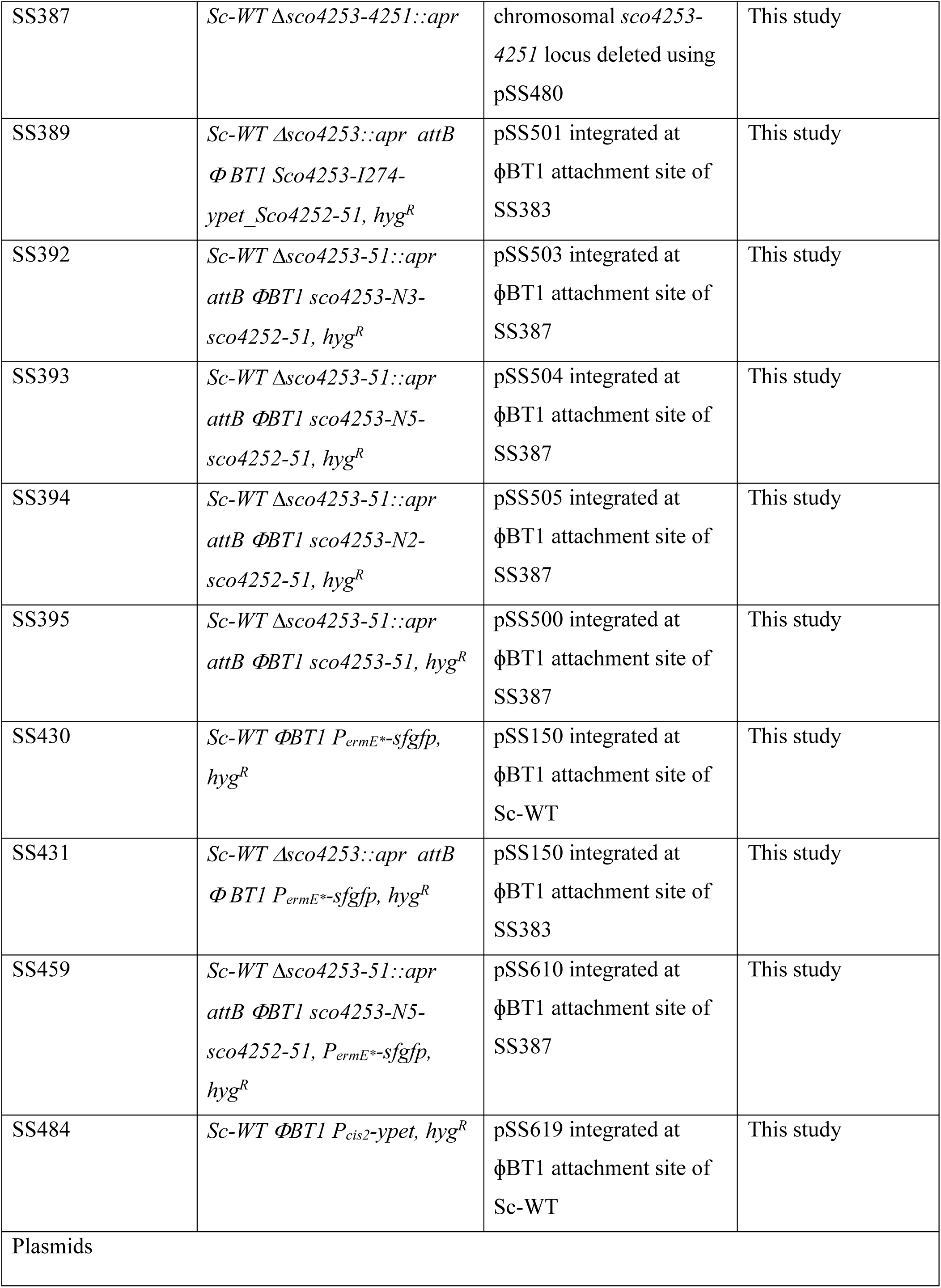
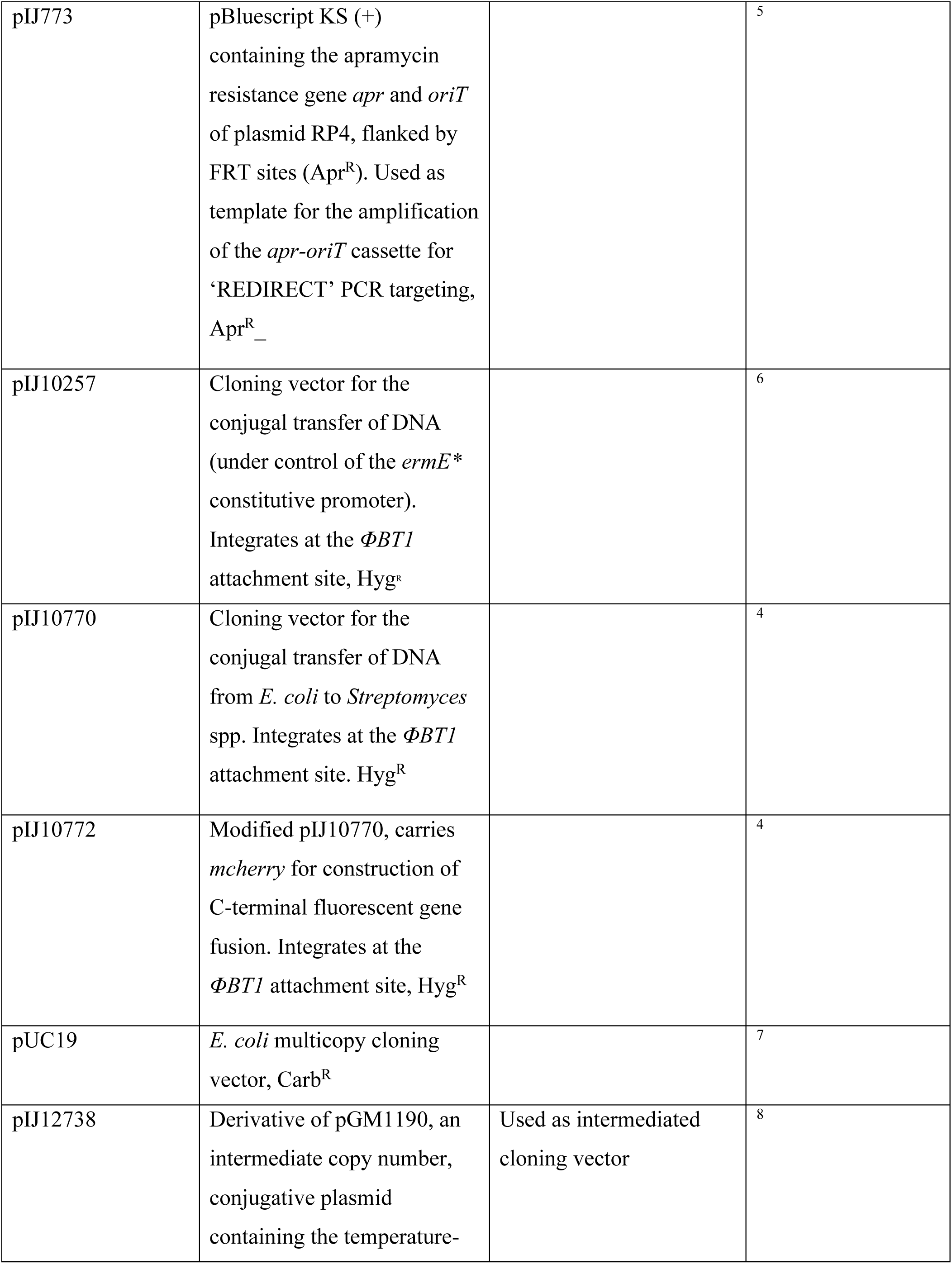
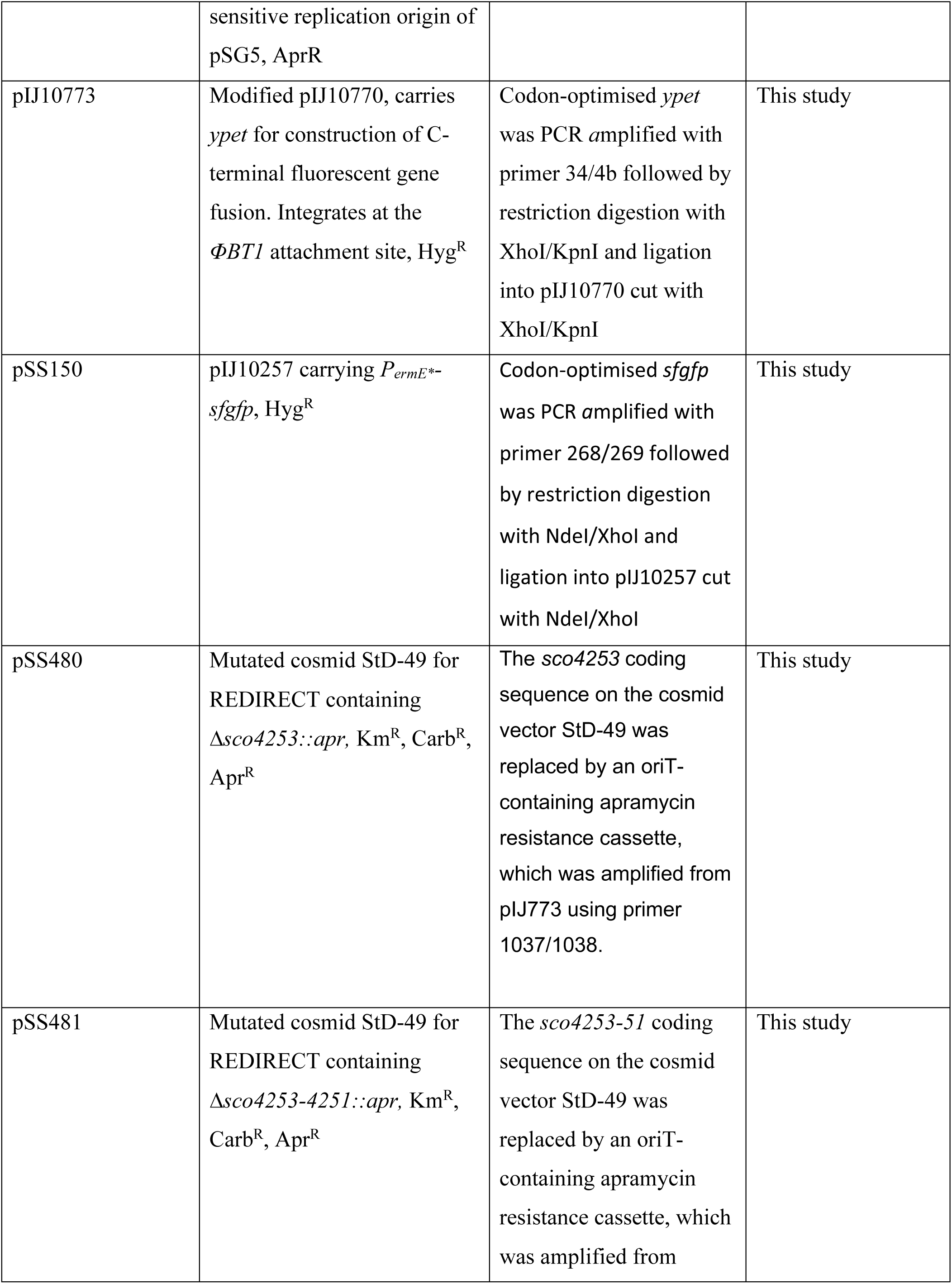
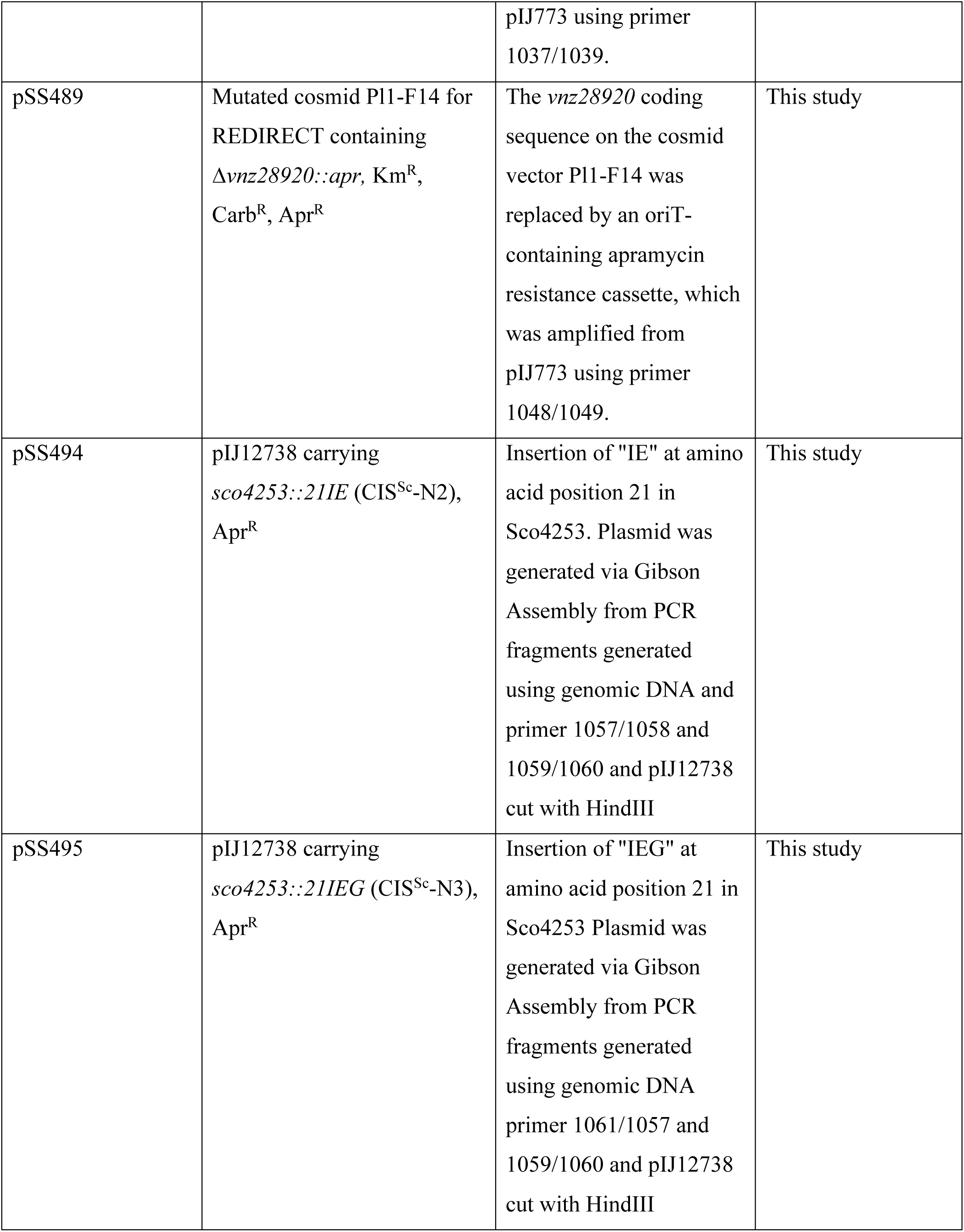
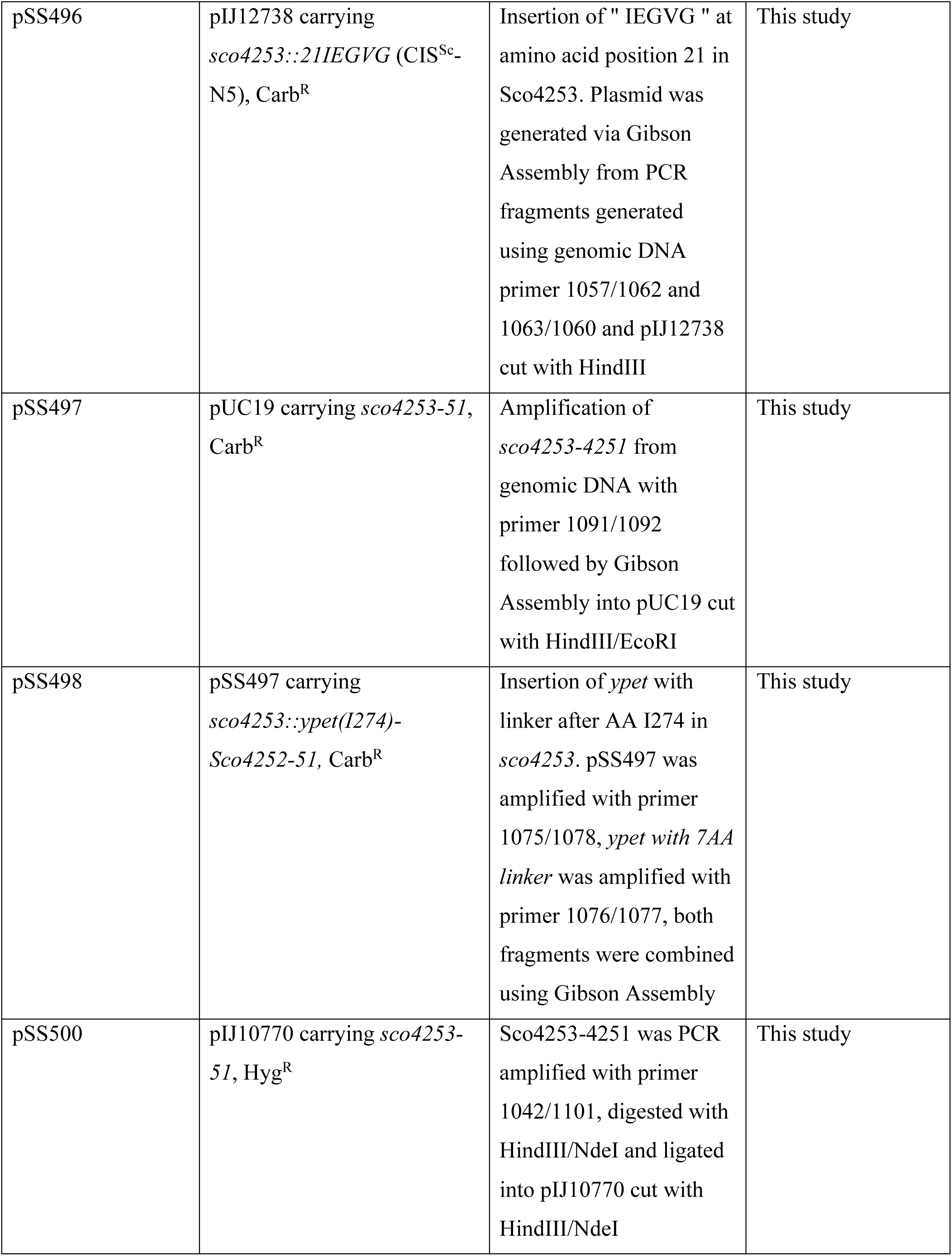
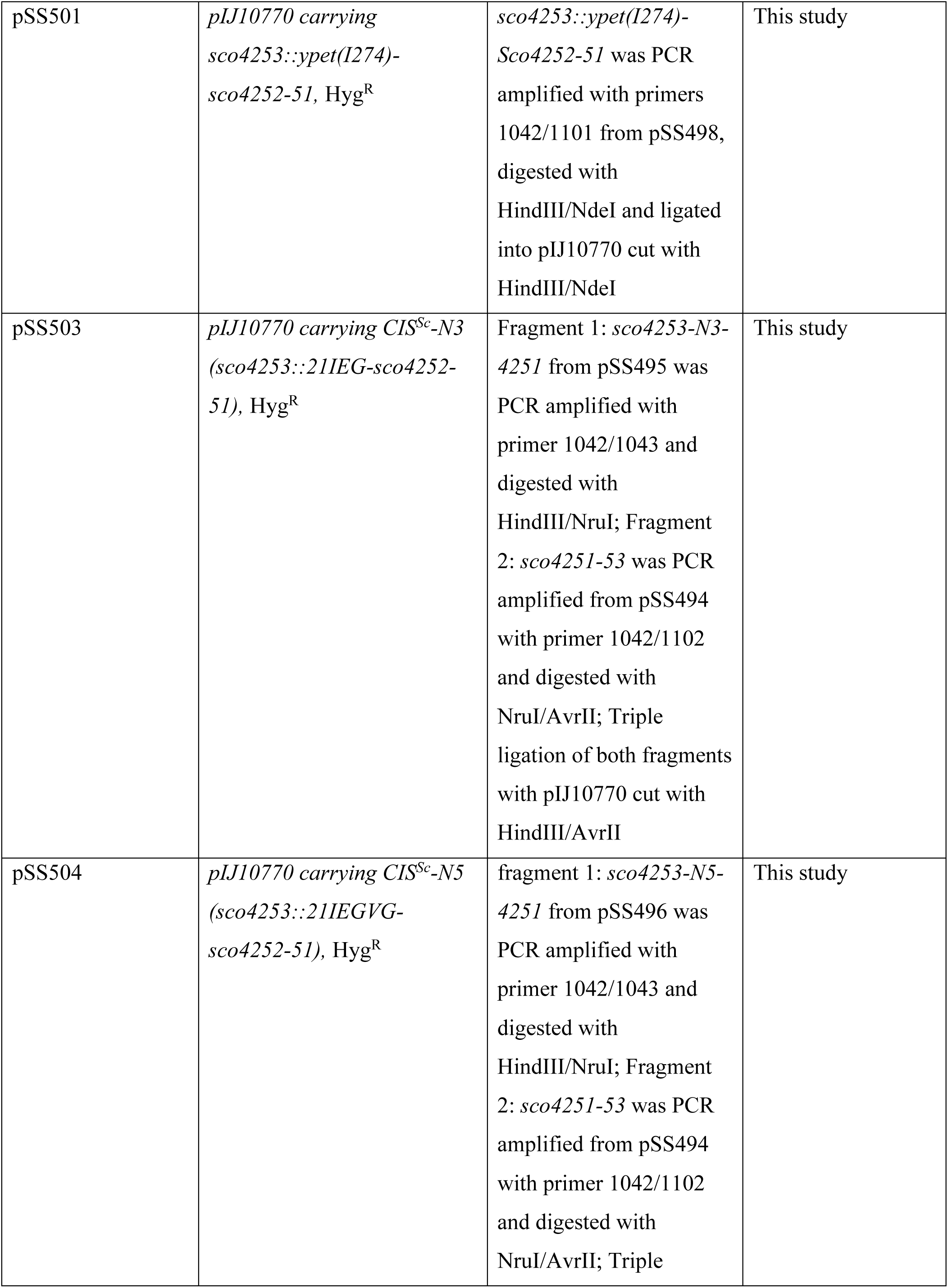
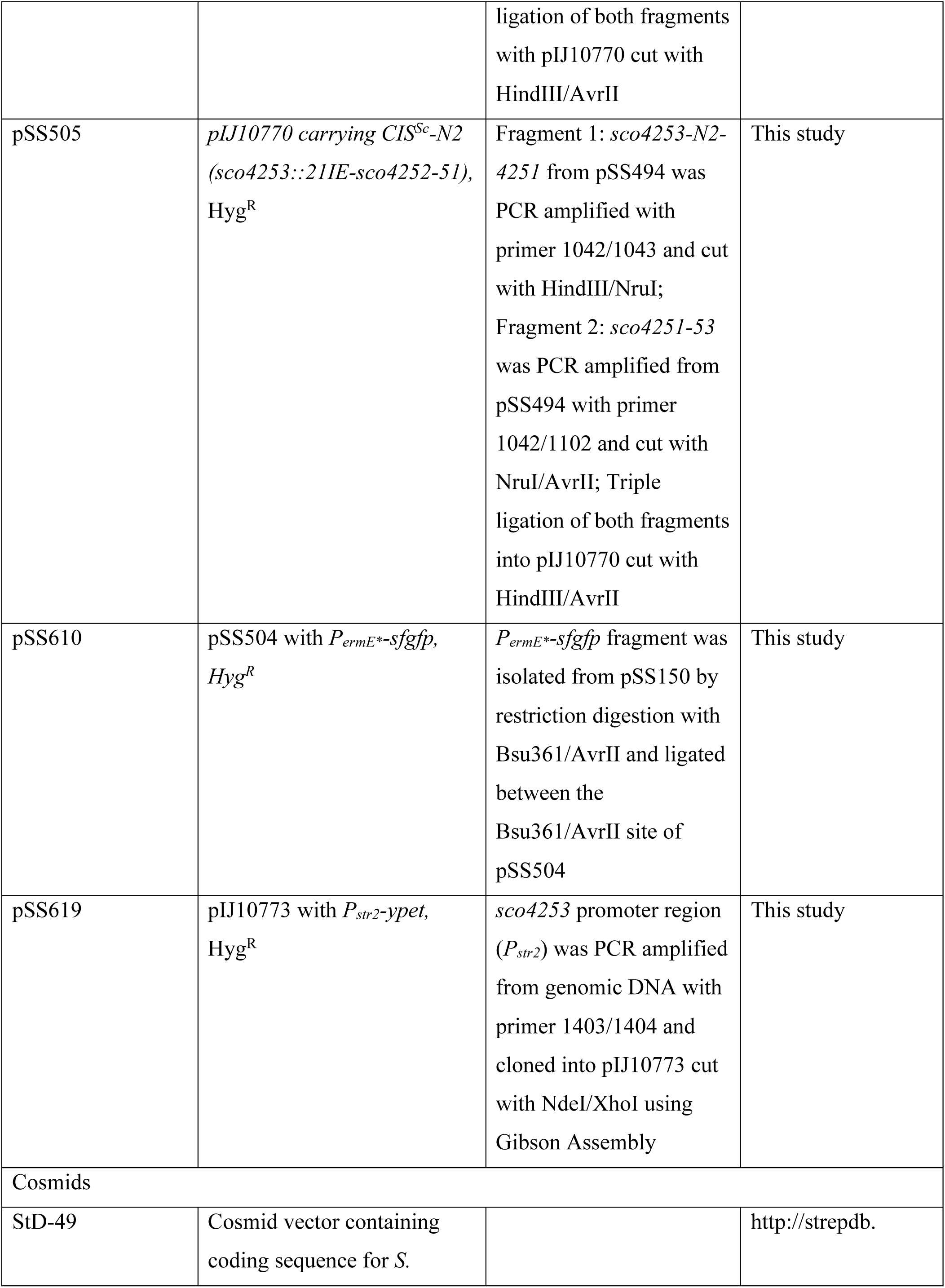
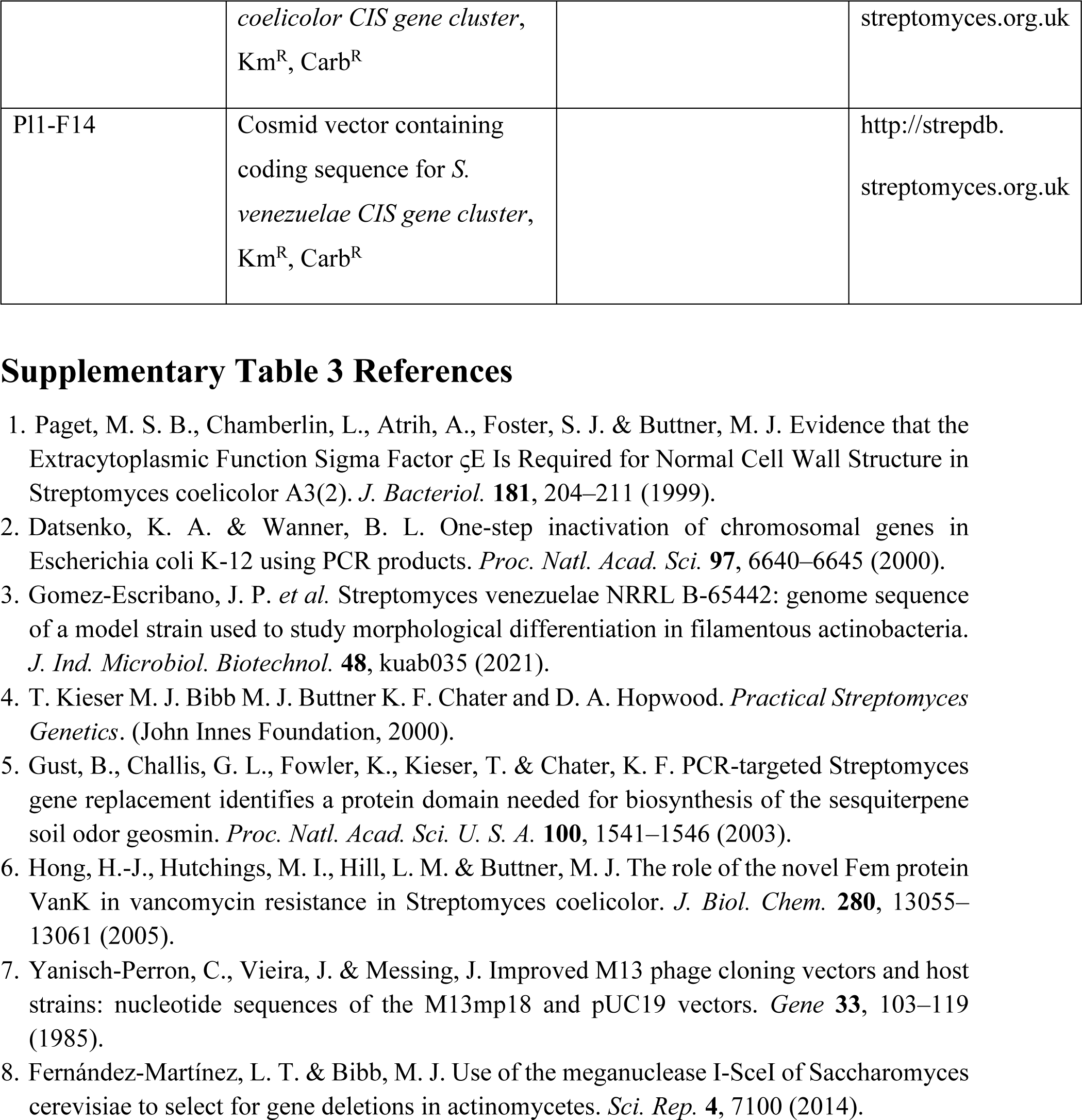
Bacterial strains, plasmids and cosmids used in this study.

**Supplementary Table 4:**
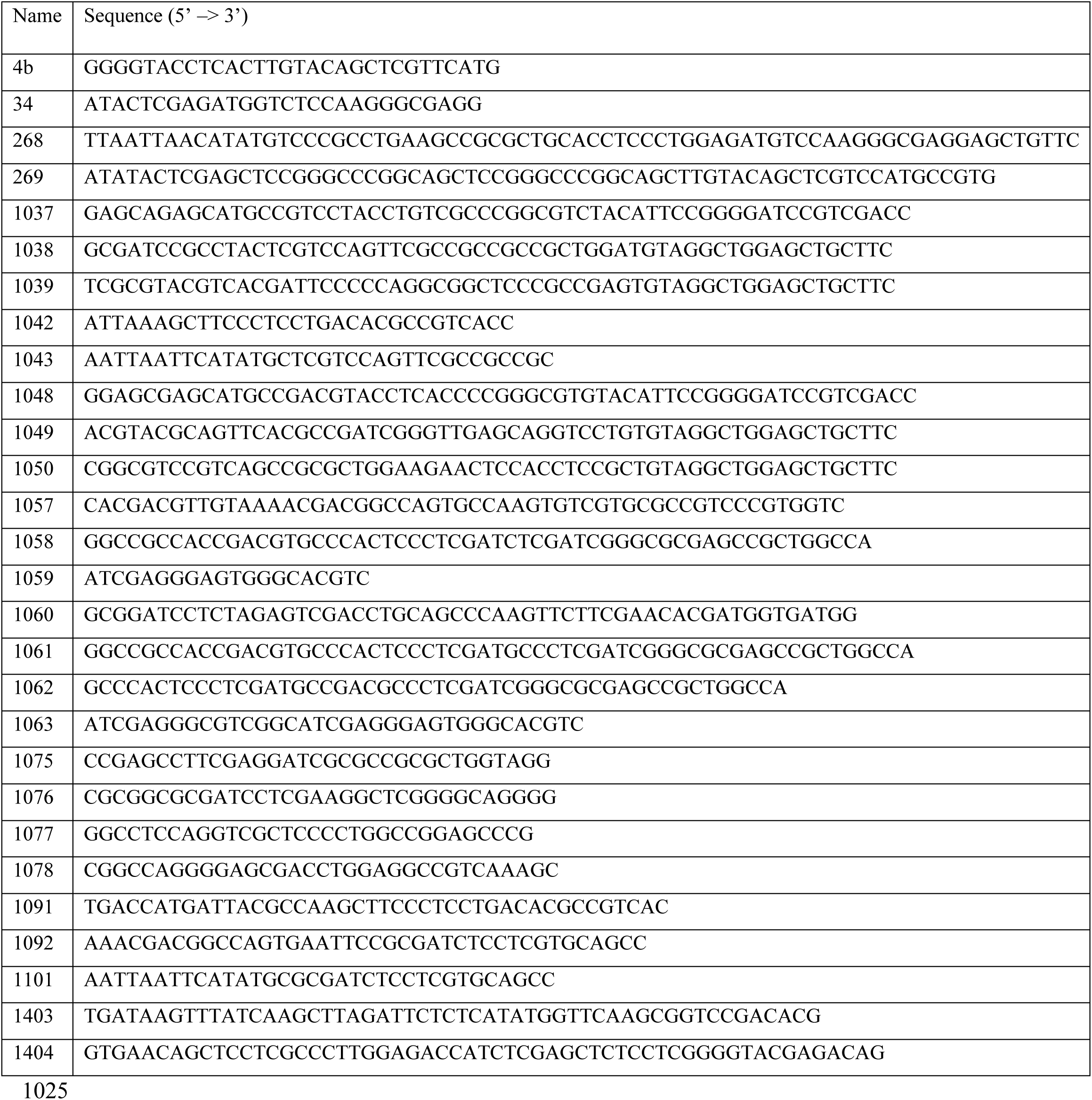
Oligonucleotides used in this study.

## References

1. Costa, T. R. D. et al. Secretion systems in Gram-negative bacteria: structural and mechanistic insights. Nat. Rev. Microbiol. 13, 343–359 (2015).

2. Veesler, D. & Cambillau, C. A Common Evolutionary Origin for Tailed-Bacteriophage Functional Modules and Bacterial Machineries. Microbiol. Mol. Biol. Rev. 75, 423–433 (2011).

3. Leiman, P. G. et al. Type VI secretion apparatus and phage tail-associated protein complexes share a common evolutionary origin. Proc. Natl. Acad. Sci. 106, 4154–4159 (2009).

4. Leiman, P. G. & Shneider, M. M. Contractile tail machines of bacteriophages. Adv. Exp. Med. Biol. 726, 93–114 (2012).

5. Taylor, N. M. I. et al. Structure of the T4 baseplate and its function in triggering sheath contraction. Nature 533, 346–352 (2016).

6. Chen, L. et al. Genome-wide Identification and Characterization of a Superfamily of Bacterial Extracellular Contractile Injection Systems. Cell Rep. 29, 511–521.e2 (2019).

7. Geller, A. M. et al. The extracellular contractile injection system is enriched in environmental microbes and associates with numerous toxins. Nat. Commun. 12, 3743 (2021).

8. Basler, M., Pilhofer, M., Henderson, G. P., Jensen, G. J. & Mekalanos, J. J. Type VI secretion requires a dynamic contractile phage tail-like structure. Nature 483, 182–186 (2012).

9. Pukatzki, S. et al. Identification of a conserved bacterial protein secretion system in Vibrio cholerae using the Dictyostelium host model system. Proc. Natl. Acad. Sci. U. S. A. 103, 1528–1533 (2006).

10. Nazarov, S. et al. Cryo-EM reconstruction of Type VI secretion system baseplate and sheath distal end. EMBO J. 37, e97103 (2018).

11. Durand, E. et al. Biogenesis and structure of a type VI secretion membrane core complex. Nature 523, 555–560 (2015).

12. Basler, M., Ho, B. T. & Mekalanos, J. J. Tit-for-tat: type VI secretion system counterattack during bacterial cell-cell interactions. Cell 152, 884–894 (2013).

13. Taylor, N. M. I., van Raaij, M. J. & Leiman, P. G. Contractile injection systems of bacteriophages and related systems. Mol. Microbiol. 108, 6–15 (2018).

14. Weiss, G. L. et al. Structure of a thylakoid-anchored contractile injection system in multicellular cyanobacteria. Nat. Microbiol. 7, 386–396 (2022).

15. Shikuma, N. J. et al. Marine tubeworm metamorphosis induced by arrays of bacterial phage tail-like structures. Science 343, 529–533 (2014).

16. Böck, D. et al. In situ architecture, function, and evolution of a contractile injection system. Science 357, 713–717 (2017).

17. Hurst, M. R. H., Beard, S. S., Jackson, T. A. & Jones, S. M. Isolation and characterization of the Serratia entomophila antifeeding prophage. FEMS Microbiol. Lett. 270, 42–48 (2007).

18. Jiang, F. et al. Cryo-EM Structure and Assembly of an Extracellular Contractile Injection System. Cell 177, 370–383.e15 (2019).

19. Xu, J. et al. Identification and structure of an extracellular contractile injection system from the marine bacterium Algoriphagus machipongonensis. Nat. Microbiol. 7, 397–410 (2022).

20. Nagakubo, T. et al. Phage tail-like nanostructures affect microbial interactions between Streptomyces and fungi. Sci. Rep. 11, 20116 (2021).

21. Hopwood, D. A. Streptomyces in Nature and Medicine: The Antibiotic Makers. (Oxford University Press, 2007).

22. Bush, M. J., Tschowri, N., Schlimpert, S., Flärdh, K. & Buttner, M. J. c-di-GMP signalling and the regulation of developmental transitions in streptomycetes. Nat. Rev. Microbiol. 13, 749–760 (2015).

23. Kudryashev, M. et al. Structure of the Type VI Secretion System Contractile Sheath. Cell 160, 952–962 (2015).

24. Ge, P. et al. Atomic structures of a bactericidal contractile nanotube in its pre- and postcontraction states. Nat. Struct. Mol. Biol. 22, 377–382 (2015).

25. Brackmann, M., Wang, J. & Basler, M. Type VI secretion system sheath inter-subunit interactions modulate its contraction. EMBO Rep. 19, 225–233 (2018).

26. Szwedziak, P. & Pilhofer, M. Bidirectional contraction of a type six secretion system. Nat. Commun. 10, 1565 (2019).

27. Wang, J. et al. Cryo-EM structure of the extended type VI secretion system sheath–tube complex. Nat. Microbiol. 2, 1507–1512 (2017).

28. P, S. & M, P. Bidirectional contraction of a type six secretion system. Nat. Commun. 10, (2019).

29. Celler, K., Koning, R. I., Willemse, J., Koster, A. J. & van Wezel, G. P. Cross-membranes orchestrate compartmentalization and morphogenesis in Streptomyces. Nat. Commun. 7, ncomms11836 (2016).

30. Wiedemann, I., Benz, R. & Sahl, H.-G. Lipid II-Mediated Pore Formation by the Peptide Antibiotic Nisin: a Black Lipid Membrane Study. J. Bacteriol. 186, 3259–3261 (2004).

31. Hesketh, A. et al. New pleiotropic effects of eliminating a rare tRNA from Streptomyces coelicolor, revealed by combined proteomic and transcriptomic analysis of liquid cultures. BMC Genomics 8, 261 (2007).

32. Bibb, M. J., Domonkos, A., Chandra, G. & Buttner, M. J. Expression of the chaplin and rodlin hydrophobic sheath proteins in Streptomyces venezuelae is controlled by σ(BldN) and a cognate anti-sigma factor, RsbN. Mol. Microbiol. 84, 1033–1049 (2012).

33. Malpartida, F. & Hopwood, D. A. Molecular cloning of the whole biosynthetic pathway of a Streptomyces antibiotic and its expression in a heterologous host. Nature 309, 462–464 (1984).

34. Tsao, S. W., Rudd, B. A., He, X. G., Chang, C. J. & Floss, H. G. Identification of a red pigment from Streptomyces coelicolor A3(2) as a mixture of prodigiosin derivatives. J. Antibiot. (Tokyo) 38, 128–131 (1985).

35. Claessen, D., Rozen, D. E., Kuipers, O. P., Søgaard-Andersen, L. & van Wezel, G. P. Bacterial solutions to multicellularity: a tale of biofilms, filaments and fruiting bodies. Nat. Rev. Microbiol. 12, 115–124 (2014).

36. Miguélez, E. M., Hardisson, C. & Manzanal, M. B. Hyphal Death during Colony Development in Streptomyces antibioticus: Morphological Evidence for the Existence of a Process of Cell Deletion in a Multicellular Prokaryote. J. Cell Biol. 145, 515–525 (1999).

37. Manteca, A., Alvarez, R., Salazar, N., Yagüe, P. & Sanchez, J. Mycelium Differentiation and Antibiotic Production in Submerged Cultures of Streptomyces coelicolor. Appl. Environ. Microbiol. 74, 3877–3886 (2008).

38. Gust, B. et al. Lambda red-mediated genetic manipulation of antibiotic-producing Streptomyces. Adv. Appl. Microbiol. 54, 107–128 (2004).

39. T. Kieser M. J. Bibb M. J. Buttner K. F. Chater and D. A. Hopwood. Practical Streptomyces Genetics. (John Innes Foundation, 2000).

40. Gust, B., Challis, G. L., Fowler, K., Kieser, T. & Chater, K. F. PCR-targeted Streptomyces gene replacement identifies a protein domain needed for biosynthesis of the sesquiterpene soil odor geosmin. Proc. Natl. Acad. Sci. U. S. A. 100, 1541–1546 (2003).

41. Edgar, R. C. MUSCLE: a multiple sequence alignment method with reduced time and space complexity. BMC Bioinformatics 5, 113 (2004).

42. Edgar, R. C. MUSCLE: multiple sequence alignment with high accuracy and high throughput. Nucleic Acids Res. 32, 1792–1797 (2004).

43. Kumar, S., Stecher, G., Li, M., Knyaz, C. & Tamura, K. MEGA X: Molecular Evolutionary Genetics Analysis across Computing Platforms. Mol. Biol. Evol. 35, 1547–1549 (2018).

44. Ohi, M., Li, Y., Cheng, Y. & Walz, T. Negative Staining and Image Classification - Powerful Tools in Modern Electron Microscopy. Biol. Proced. Online 6, 23–34 (2004).

45. Bush, M. J., Gallagher, K. A., Chandra, G., Findlay, K. C. & Schlimpert, S. Hyphal compartmentalization and sporulation in Streptomyces require the conserved cell division protein SepX. Nat. Commun. 13, 71 (2022).

46. Bush, M. J., Bibb, M. J., Chandra, G., Findlay, K. C. & Buttner, M. J. Genes Required for Aerial Growth, Cell Division, and Chromosome Segregation Are Targets of WhiA before Sporulation in Streptomyces venezuelae. mBio 4, e00684–13 (2013).

47. Schindelin, J. et al. Fiji - an Open Source platform for biological image analysis. Nat. Methods 9, 10.1038/nmeth.2019 (2012).

48. Medeiros, J. M. et al. Robust workflow and instrumentation for cryo-focused ion beam milling of samples for electron cryotomography. Ultramicroscopy 190, 1–11 (2018).

49. Weiss, G. L., Medeiros, J. M. & Pilhofer, M. In Situ Imaging of Bacterial Secretion Systems by Electron Cryotomography. Methods Mol. Biol. Clifton NJ 1615, 353–375 (2017).

50. Mastronarde, D. N. Automated electron microscope tomography using robust prediction of specimen movements. J. Struct. Biol. 152, 36–51 (2005).

51. Kremer, J. R., Mastronarde, D. N. & McIntosh, J. R. Computer visualization of three-dimensional image data using IMOD. J. Struct. Biol. 116, 71–76 (1996).

52. Tegunov, D. & Cramer, P. Real-time cryo-electron microscopy data preprocessing with Warp. Nat. Methods 16, 1146–1152 (2019).

53. Zheng, S. Q. et al. MotionCor2: anisotropic correction of beam-induced motion for improved cryo-electron microscopy. Nat. Methods 14, 331–332 (2017).

54. Zivanov, J. et al. New tools for automated high-resolution cryo-EM structure determination in RELION-3. eLife 7, e42166 (2018).

55. He, S. & Scheres, S. H. W. Helical reconstruction in RELION. J. Struct. Biol. 198, 163– 176 (2017).

56. Rosenthal, P. B. & Henderson, R. Optimal determination of particle orientation, absolute hand, and contrast loss in single-particle electron cryomicroscopy. J. Mol. Biol. 333, 721– 745 (2003).

57. Ef, P. et al. UCSF Chimera--a visualization system for exploratory research and analysis. J. Comput. Chem. 25, (2004).

58. Emsley, P., Lohkamp, B., Scott, W. G. & Cowtan, K. Features and development of Coot. Acta Crystallogr. D Biol. Crystallogr. 66, 486–501 (2010).

59. Song, Y. et al. High-resolution comparative modeling with RosettaCM. Struct. Lond. Engl. 1993 21, 1735–1742 (2013).

60. Adams, P. D. et al. PHENIX: a comprehensive Python-based system for macromolecular structure solution. Acta Crystallogr. D Biol. Crystallogr. 66, 213–221 (2010).

61. Goddard, T. D. et al. UCSF ChimeraX: Meeting modern challenges in visualization and analysis. Protein Sci. Publ. Protein Soc. 27, 14–25 (2018).

62. Schorb, M., Haberbosch, I., Hagen, W. J. H., Schwab, Y. & Mastronarde, D. N. Software tools for automated transmission electron microscopy. Nat. Methods 16, 471–477 (2019).

63. Hurst, M. R. H., Glare, T. R. & Jackson, T. A. Cloning Serratia entomophila Antifeeding Genes—a Putative Defective Prophage Active against the Grass Grub Costelytra zealandica. J. Bacteriol. 186, 5116–5128 (2004).

64. Zhang, Y., Wang, L., Zhang, S., Yang, H. & Tan, H. hmgA, transcriptionally activated by HpdA, influences the biosynthesis of actinorhodin in Streptomyces coelicolor. FEMS Microbiol. Lett. 280, 219–225 (2008).

